# Relative contribution of gonads and sex chromosomes to sex differences in cell-type gene expression in the mouse medial septum and sex-biased disease risk

**DOI:** 10.64898/2025.12.08.693050

**Authors:** Juliana Shin, Montgomery Blencowe, Caden McQuillen, Haley Hrncir, Xuqi Chen, Graciel Diamante, In Sook Ahn, Guanglin Zhang, Arthur P. Arnold, Allan Mackenzie-Graham, Jason Lerch, Armin Raznahan, Xia Yang

## Abstract

**Background:** While sex differences in the brain have traditionally been attributed to gonadal hormones, emerging evidence points to regulation by sex chromosomes. This study aims to differentiate the influence of gonads versus sex chromosomes on cellular gene expression in the mouse medial septum (MS), a critically understudied brain region.

**Methods:** Using single nucleus RNA-sequencing and the Sex Chromosome Trisomy mouse model, we (1) quantified sex differences in cellular gene expression and (2) isolated sex-biasing effects by identifying perturbed cell types, differentially expressed genes, biological pathways, and gene networks, which were integrated with GWAS data to explore links with sex-biased human phenotypes.

**Results:** Our analysis revealed that volumetric sex differences in the MS are mirrored by widespread transcriptomic changes across cell types. Critically, genetic effects displayed elevated relevance compared to sex hormones in driving sex-biased gene expression. These effects converge to regulate synaptic/neuronal development, transcriptional regulation, and cellular metabolism. Sex chromosome-associated DEGs were enriched for various human disorders, suggesting a cellular and mechanistic basis for their sex-biased patterns.

**Conclusions:** Our findings challenge the classical gonad-centric views of sexual differentiation, as the MS displays sex-biased transcriptional regulation driven by sex chromosome-associated effects that are highly relevant for human health.

## Introduction

Pronounced sex differences exist throughout the mammalian brain and manifest as robust phenotypic and structural differences between males and females. Given these well-established sex differences in diverse human behaviors and mental health outcomes, there has been growing interest in interrogating patterns of sex-differentiated brain development and function. One approach, in vivo neuroimaging methods, has proven successful in profiling both regional sex differences in brain anatomy within large cohorts of typically developing individuals^1,2^, as well as regional variation in brain anatomy caused by gonadal hormones^3^ and sex chromosome dosage^4^. Parallel neuroimaging studies in mice reveal partially overlapping sex differences in brain volume to those seen in humans^5^, and by exploiting the greater accessibility to targeted experimental approaches in mice, studies have also confirmed a causal role for gonadal type and sex chromosome dosage in influencing regional brain anatomy^6,7^.

Because humans and mice share several aspects of sex-biased biology, detailed dissection of sex differences afforded by murine studies provide important leads on potential causal processes that prove difficult to parse in humans. For example, experimental studies of canonical foci of male-biased brain volume in mice and humans, such as the medial amygdala and bed nucleus of the stria terminalis, point to a dominant influence of testicular hormone exposure on brain development^3,8–13^. To date, however, most studies attempting to identify the underpinnings of sex-biased brain anatomy in mice and humans has relied on bulk gene expression data, which lacks both cellular resolution as well as the ability to separate the effects of distinct sex-biasing factors^14–19^. We therefore lack an understanding of not only molecular sex differences at single cell scale in foci of sex-biased anatomy, but also the potential causal roles of gonads and sex chromosomes, especially within the more recently discovered regions of female-biased volume in the mouse brain^20^. One of such female-biased regions is the medial septum (MS) that also appears to be volumetrically sensitive to sex chromosome, but not gonadal, effects^4,6,21^. Notably, removal of gonadal hormones prior to puberty did not affect the XX > XY pattern observed in the MS^22^.

The MS is a component of the basal forebrain that projects to and forms reciprocal connections with the hippocampus via a complex regulatory network featuring GABAergic, glutamatergic, and cholinergic neurons. The MS is responsible for the regulation of theta rhythmogenesis, a form of neural oscillation essential for various cognitive functions that notably often exhibit sex differences, including spatial navigation, memory consolidation, and attention maintenance^23^. Beyond its role in the septo-hippocampal circuit and evidence of sex differences in neuroanatomy, the activity of the MS in mice also appears to vary as a function of sex and sex-biasing factors. For example, neurodevelopmental studies have identified stable sex differences in the pattern of neurogenesis of cholinergic neurons of the MS, with peak neurogenesis occurring earlier in male than female mice^24^. Furthermore, gonadal hormones, specifically estrogens but not testosterone, directly regulate cholinergic neurons in the forebrain cholinergic system via modulation of neurotrophic receptors^25,26^. Studies have also identified greater male vulnerability to impairments in memory formation mediated by circulating corticotropin releasing factor (CRF) in the MS that is independent of circulating gonadal hormones and alters MS-mediated memory formation^27^. Cumulatively, these findings raise the yet untested hypothesis that the cellular gene programming and organization underlying the MS may be sensitive to both gonadal and sex chromosome complement/dosage effects. These effects may disturb proportions and gene expression signatures of the major MS cell types including neuronal subtypes (GABAergic, Glutamatergic, and Cholinergic) and various non-neuronal cell types. Such an understanding of the MS and its full functions are currently lacking, as the MS is a critically understudied brain region.

Here, we seek to address these gaps in knowledge through single nucleus RNA-sequencing (snRNA-seq) analyses of gonadal hormone and sex chromosome dosage effects in MS using the unique explanatory power of the Sex Chromosome Trisomy (SCT) mouse model^28,29^. In these mice, the *Sry* gene is deleted from the Y chromosome but present as a transgene on chromosome 3, so that the gonadal type (testes, denoted as T, vs ovaries, denoted as O) of these mice is independent of their sex chromosome complement. The mothers of the SCT cross are XXY (with the Y chromosome lacking *Sry*), who produce two types of eggs, X or XY. With four separate chromosome groups (XX, XY, XXY, XYY), the model features 8 distinct genotype cohorts (XXO, XXT, XYO, XYT, XXYO, XXYT, XYYO, XYYT) (**Supp. Fig 1**). Thus, snRNA-seq in the SCT model will not only assay “typical” sex differences (represented by the contrast between XYT and XXO groups), but also the dissociable effects of gonadal type (O vs T), denoted as “gonadal effects” (GEs), and sex chromosome complement effects (XX vs XY), denoted as “sex chromosome effects” (SCEs) by using the “Four Core Genotypes” (FCG) subset of the SCT model (XXO, XXT, XYO, XYT)^30^.

Moreover, the SCT model can refine potential causal bases for observed SCEs by separately estimating the effects of variation in X- and Y-chromosome dosage (XCD, comparing XY vs XXY; and YCD, comparing XX vs. XXY and XY vs XYY). These sex chromosome aneuploid groups can simultaneously shed light on cellular mechanisms in human sex chromosome aneuploidy syndromes such as Turner and Klinefelter syndrome. Therefore, the use of the SCT model coupled with snRNAseq can dissect the unique contributions of all sex factors, including FCG effects (GE: O vs T; SCE: XX vs XY; GExSCE) and SCT effects (XCD: 2 vs 1 copies of X; YCD: 2 vs 1 vs 0 copies of Y) at the cell type and transcriptomic level by analyzing sex driven effects at single nuclei resolution. We use our study design to interrogate how sex factors perturb cellular composition, gene expression, biological pathways, gene regulatory networks (GRNs), and potential disease association in a critically understudied brain region.

## Methods

### SCT mice breeding and genotyping

Outbred MF1 mice of the Sex Chromosome Trisomy model were used (ref PMID: 23926958). Like Four Core Genotypes mice (FCG, Jackson strain 039108) (PMID: 32980399), these mice harbor a Y chromosome with the dl1Rlb allele deleted for *Sry*, designated Y^─^, and an autosomal transgene for *Sry* inserted into Chromosome 3. Gonadal female XXY^─^ mice were crossed with XY^─^ (*Sry*+) gonadal males to produce 8 types of offspring: XX, XY^─^, XXY^─^ and XY^─^Y^─^, each genotype either lacking *Sry* and possessing ovaries, or possessing *Sry* and testes. All mice derived from mice containing an identical X chromosome that Paul Burgoyne produced by crossing an XO mother with her XY son and then propagating the individual X chromosome throughout the colony. Thus, differences caused by dosage of the X chromosome are not the result of allelic differences among X chromosomes. The MF1 strains of FCG and SCT mice do not possess the translocation of X genes to the Y^─^ chromosome found in some lines of C57BL/6J FCG mice^31^. Mouse genotypes were determined using genomic PCR to detect the Y chromosome and *Sry* transgene^32^. Fluorescent in situ hybridization on interphase lymphocytes, using probes for the X and Y chromosomes, was used to detect the number of X and Y chromosomes (Kreatek kit KI-30505, Leica Biosysystems, USA). A visual schematic of the SCT mouse model is shown in **Supp Figure 1**.

### MS dissection, nuclei isolation, and snRNA-seq library preparation

Mice of an average of P92 days were anesthetized with isoflurane and sacrificed. Brains were sectioned in an acrylic coronal mouse brain matrix to isolate a 1mm section anterior of the decussation of the anterior commissure containing the MS. The MS, as defined in The Mouse Brain in Stereotaxic Coordinates^33^, was subsequently dissected with a scalpel and immediately flash frozen in liquid nitrogen.

Nuclei were isolated from the MS samples (n=3 independent mice/group for a total of 24 samples across 8 groups) using the Minute Detergent-Free Single Nuclei Isolation Kit following the manufacturer’s experimental procedure (Invent Biotechnologies, INC, Plymouth, MN, USA). Single-nuclei suspensions were then used to generate Gel Bead-in-emulsions (GEMs) and sequencing libraries following the 10x 3′ single-cell RNA-seq V3.1 protocol (10x Genomics, Pleasanton, California, USA). The concentration of the libraries was determined using the Qubit Fluorometric Quantitation method (Thermofisher Scientific, Waltham, MA, USA), and quality was assessed using the Agilent TapeStation system (Agilent, Santa Clara, CA, USA). The pooled libraries were then sequenced on the Novaseq S4 2×100 (Illumina, San Diego, CA, USA) in the UCLA Broad Stem Cell Research Center at ~35k reads/cell.

### snRNA-seq pre-processing and quality control

Raw base call files (BCL) were demultiplexed and converted to FASTQ format, and FASTQ files were mapped to the mm10 2020a reference genome using 10x Genomics’ Cell Ranger software (v6.0.2). Intronic reads were included as snRNA-seq profiles nuclear transcripts that include introns. To remove signals derived from ambient RNA contamination, Cell Ranger output files were processed using the CellBender package (v0.3.0)^34^. Corrected individual sample-wise gene expression matrices were loaded and combined using Seurat (v4.1.1)^35^. Nuclei with <200 and >7,500 UMIs, feature counts < 500 or >35,000, and >3% mitochondrial percentage were filtered out. Remaining nuclei were annotated as predicted singlets or doublets using DoubletFinder (v2.0.4)^36^. Quality control plots are shown in **Supp Figure 2**.

### Data normalization, batch correction, and cell clustering

The gene expression matrix for individual samples was normalized using Seurat’s “SCTransform” function v2 and then subject to principal component analysis (PCA) using Seurat’s “RunPCA” function. To correct for batch effects, Seurat’s “RunHarmony” function was used, with the grouping variable set to “Sequencing Date”. To cluster nuclei with similar expression patterns, the K-Nearest Neighbors and Louvian algorithms were used via Seurat’s “FindNeighbors” and “FindCluster” functions. Lastly, Uniform Manifold Approximation Projection (UMAP) was performed to visualize clusters in two-dimensional space using the top 30 dimensions. Clusters with high doublet abundance were removed, leaving an average of 5863 nuclei per sample for downstream analysis.

### Cell Type Annotations

Clusters were annotated based on expression of literature-derived canonical cell type markers as well as MS-specific neuronal markers^37–39^. Annotations were validated using the Allen Institute’s MapMyCells tool (RRID:SCR_024672).

### Genotype Validation Using snRNAseq Data

To confirm the genotype of each sample initially genotyped using interphase FISH, we assessed *Xist* and *Uty* expression post-sequencing. We identified 4 mis-genotyped samples and reassigned genotypes for the 8 groups, including XYT (n = 5), XYO (n = 2), XXT (n = 3), XXO (n = 4), XYYT (n = 3), XYYO (n = 3), XXYT (n = 1), and XXYO (n = 3) (**Supp Figure 3**). This revised genotyping led to the following sample sizes: typically developing sex (n=5 XYT vs n=4 XXO), gonadal (n=12 T vs 12 O), sex chromosome complement (n=7 XX vs n=7 XY), X dosage (n=13 with one X vs n=11 with two X), Y dosage (n=7 with 0 Y vs n=11 with one Y vs n=6 with 2 Y). Of note, these sample sizes are larger than typically used in single cell studies in mice.

### Cell Type Proportion Analysis

To assess differences in cell type composition between groups, cell type proportions were calculated for each sample individually as the total number of nuclei for each cell type divided by the total number of nuclei in each sample. Proportions were similarly calculated by genotype, gonad type, sex chromosome complement, and trisomy status. Significant differences in cell type compositions between groups were tested for using the propeller method within the speckle R package (v1.6.0)^40^.

### Generation of metacells and Identification of Differentially Expressed Genes (DEGs)

Recent benchmarking studies support the use of metacells over single cell and pseudobulk approaches in DEG identification^41^. The metacell approach groups transcriptionally similar cells into a single representative cell state, termed a “metacell”, to construct multiple metacells of a given cell type within a given sample. This approach better preserves biological variation while also reducing data sparsity and biases stemming from pseudoreplication and has been successfully applied in diverse settings to determine robust and biologically meaningful DEGs^42–46^. Metacells were generated within sample and cell type using the SuperCell package and default parameters^47^. Final metacell proportions per sample and cell type reflect original cell type abundances (**Supp Fig 4**).

To determine the contributions of the typical sex difference and individual sex factors to gene expression in each cell type, gene-wise linear models were fitted across all relevant samples using the Limma R package (v3.58.1)^48^ to calculate: (1) the typical sex difference, (2) gonadal effects (GEs), (3) sex chromosomal effects (SCEs), (4) interactions between GE and SCEs (GExSCE), (5) X chromosomal dosage (XCDs), and (6-7) Y chromosomal dosages (YCD1: 0 vs 1 Y; YCD2: 1 vs 2 Y). All linear models included sample sequencing date as a covariate to account for technical batch effects. A summary of each model’s parameters and relevant samples are included in **Supp Table 1**.

For the typical sex difference, the linear model used was:

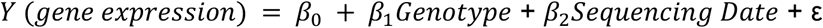

with the specific contrast set to *XXOvaries - XYTestes*.

The second linear model was used to infer “FCG Effects” (i.e., those typically measured in the four FCG genotypes which comprise half of the SCT model), which are gonadal and sex chromosome effects as well as their interactions used mice of the XYT, XYO, XXT, and XYO genotypes:

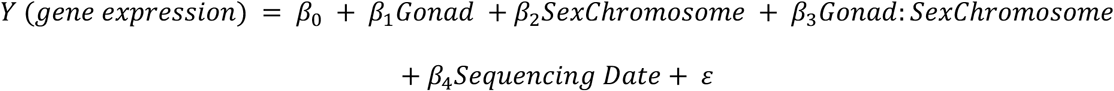

Separate models were used to infer the XCD and YCD effects with relevant genotypes only. In order to infer the effects of X/Y chromosome aneuploidy, all models featured a “base Genotype” term that summarized the effects of the Genotype in the absence of the added sex chromosome. For example, for a XXYT sample, its base Genotype term for the XCD term was set to XYT, while its base Genotype term for the YCD term was set to XXT. Moreover, given the dosage-sensitive nature of Y chromosome genes^49^, two models were used to isolate the influence of 1 vs. 0 additional Y chromosomes (YCD1) or by 2 vs. 1 additional Y chromosomes (YCD2).

The linear models used were as follows:

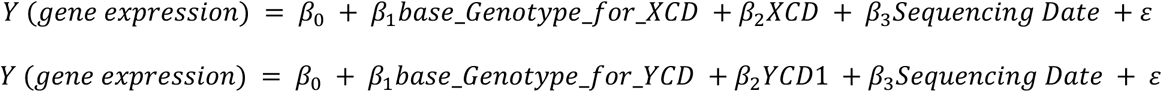

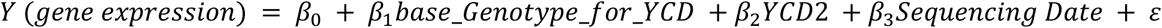

Differential gene expression was calculated from the above models and false discovery rate (FDR) was calculated using the Benjamini-Hochberg (BH) method. DEGs with FDR < 0.05 and log2 fold change (log2FC) > 0.1 were considered significantly differentially expressed.

### Gene log2FC correlation and regression analysis across sex-biasing factors

To assess the contribution of individual sex factors to typical sex differences and infer similarity between the transcriptomic effects of individual sex factors, log2FC values across all measured genes were isolated for each cell type and sex effect pair. For each cell type, the log2FC for the typical sex difference effect was then correlated with the log2FC for all other sex factors using the *cor* function in R and the Pearson method. A linear regression model was also fit using the typical sex difference log2FC value as a response variable, and the other sex effect log2FC values as explanatory variables for each cell type. Positive or negative correlation values from Pearson correlation and beta values from linear models were plotted to understand how individual sex factors relate to and contribute to typical sex differences.

### Clustering analysis of DEGs across cell types and sex factors and Pathway Enrichment Analysis of DEG clusters

Due to the large numbers of DEG sets across cell types and sex factors, we first identified DEG sets sharing similar patterns before interpreting their implicated biological pathways. The log2FC values of DEGs were isolated for each cell type and sex effect pair and correlated with each other using the Pearson method. The cell type and sex effects DEG sets were grouped using hierarchical clustering based on log2FC correlation values. Distinct clusters were subsequently defined by cutting the resulting dendrogram tree at a height that best constructed biologically interpretable gene clusters (h = 2.6). Significant DEGs present in at least 20% DEG sets within a cluster were then used as input for pathway enrichment for biological interpretation using the clusterProfiler package (v4.10.0) and gene sets from the GO Biological Process (GO:BP) database^50^. Pathways at FDR<5% were deemed significantly overrepresented. Redundant pathways that featured highly overlapping genes were identified using the Jaccard index coefficient and non-redundant representative pathways were selectively presented.

### Gene Regulatory Network (GRN) Construction

Global GRNs were constructed by SCING using cell type-specific gene count matrices^51^. Gradient boosting regressors were trained on each gene to form directed edges from potential upstream regulators. Following network inference, weighted Key Driver Analysis (wKDA) module from the Mergeomics R package was used to overlay cell type-specific sex factor DEG sets onto constructed networks, allowing for identification of key drivers whose neighborhood subnetworks were significantly enriched for DEGs at FDR<5%^52^. For each cell type, the top 5 key drivers for each sex effect were selected for visualization.

### Disease Enrichment Analysis

To examine the potential disease relevance of cell type-specific sex factor DEGs to human disease, a total of 98 summary statistics of human GWAS for a range of neuropsychiatric, metabolic, and locomotory traits and diseases were collected (**Supp Table 27**). GWAS SNPs were linked to genes via 50kb distance mapping and corrected for linkage disequilibrium using the Marker Dependency Filtering (MDF) module of Mergeomics. Significant sex factor DEGs from our mouse SCT model were converted to human orthologs. The Marker Set Enrichment Analysis (MSEA) function from the Mergeomics R package was applied to determine if sex factor DEGs show stronger GWAS association than would be expected by random chance by comparing disease association p-values of SNPs mapped to our DEG sets to those of the SNPs mapped to 10000 sets of random genes using a χ^2^-like statistic. Permutation-based p-values were calculated and the BH method was used to correct for muse traits. Significance was set to FDR<5%.

## Results

### snRNA-seq reveals cellular landscape of the MS in SCT mice

Cells across all 8 SCT groups (n = 24) were clustered based on transcriptomic similarity and projected to a 2D space using UMAP. After quality control and filtering, a total of 150,469 nuclei were included in our analysis, featuring 31789 nuclei from the XYT group, 16043 from XYO, 15601 from XXT, 18646 from XXO, 24008 from XYYT, 17846 from XYYO, 10691 from XXYT, and 15845 from XXYO. We identified a total of 10 major neuronal and non-neuronal cell types, including inhibitory immature neurons (Inh-IMN), astrocytes, ependymal cells, microglia, oligodendrocytes (Oligo), oligodendrocyte precursor cells (OPC), vascular cells, and three neuronal subtypes known to populate the MS, glutamatergic (Glut), GABAergic (GABA), and cholinergic (Chol) neurons (**Figure 2A**). Cluster marker genes aligned with well-documented cell markers (**Figure 2B,C**). A small cluster of cells that showed moderate expression of multiple neuronal markers were annotated as “Miscellaneous neurons” (Misc-NT) and removed from downstream data analysis.

**Figure 1.**
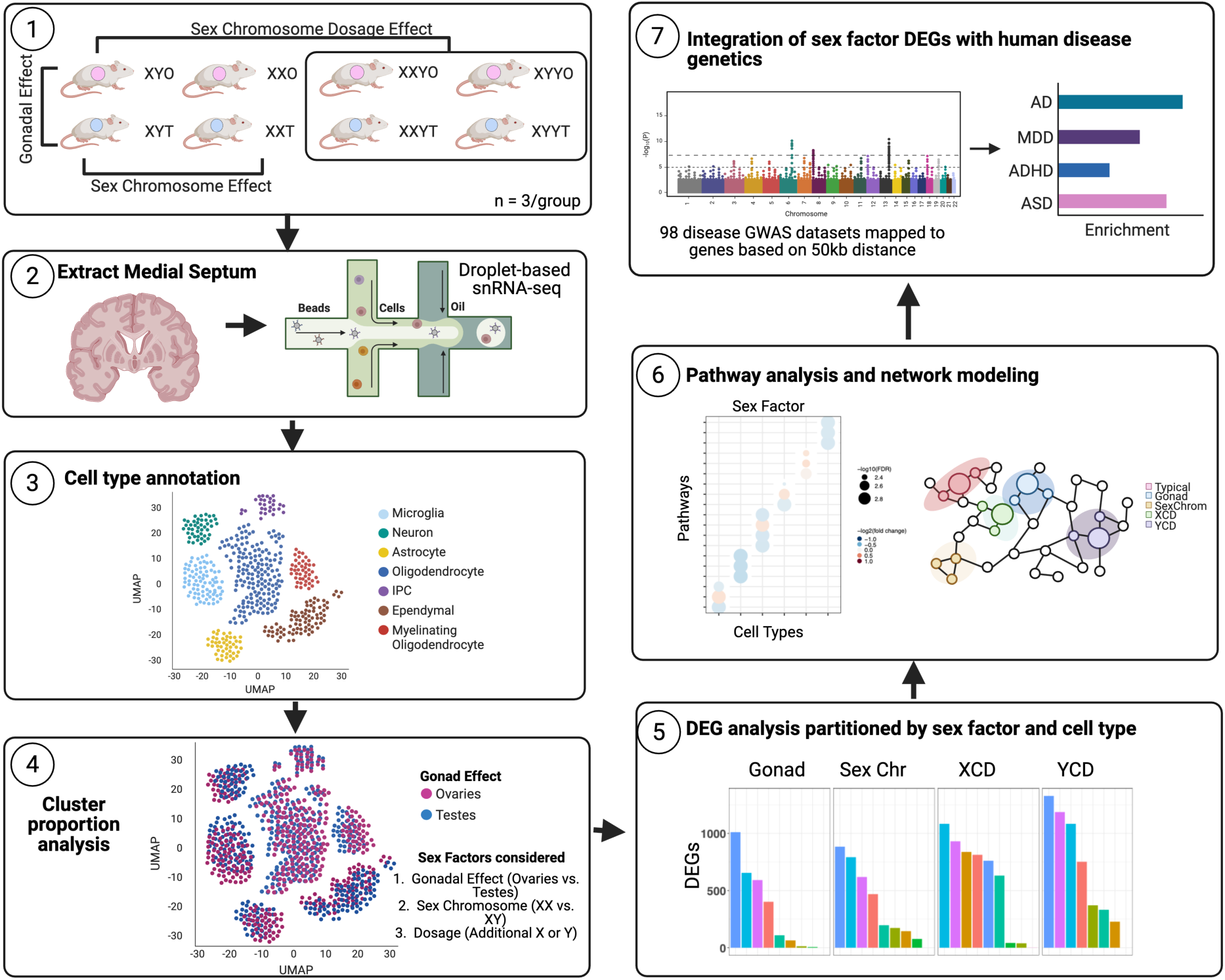
**Experimental and analysis overview.** (1) Schematic of SCT mouse model, with contrasts for each assessed sex-biasing effect outlined. (2) Dissection of the medial septum tissue for RNA isolation and snRNA-seq. (3) Cell annotation based on marker gene expression. (4) Differential analysis of cell type/cluster proportions. (5) Differentially expressed genes influenced by individual sex-biasing factors were isolated via *limma*. (6) DEGs were analyzed for enrichment of functional pathways and used for network analysis to construct cell type-specific GRNs. (7) Relevance of DEGs with human disease genetics were assessed via integration with human genome-wide association studies (GWAS) for 98 diseases using MSEA.

**Figure 2.**
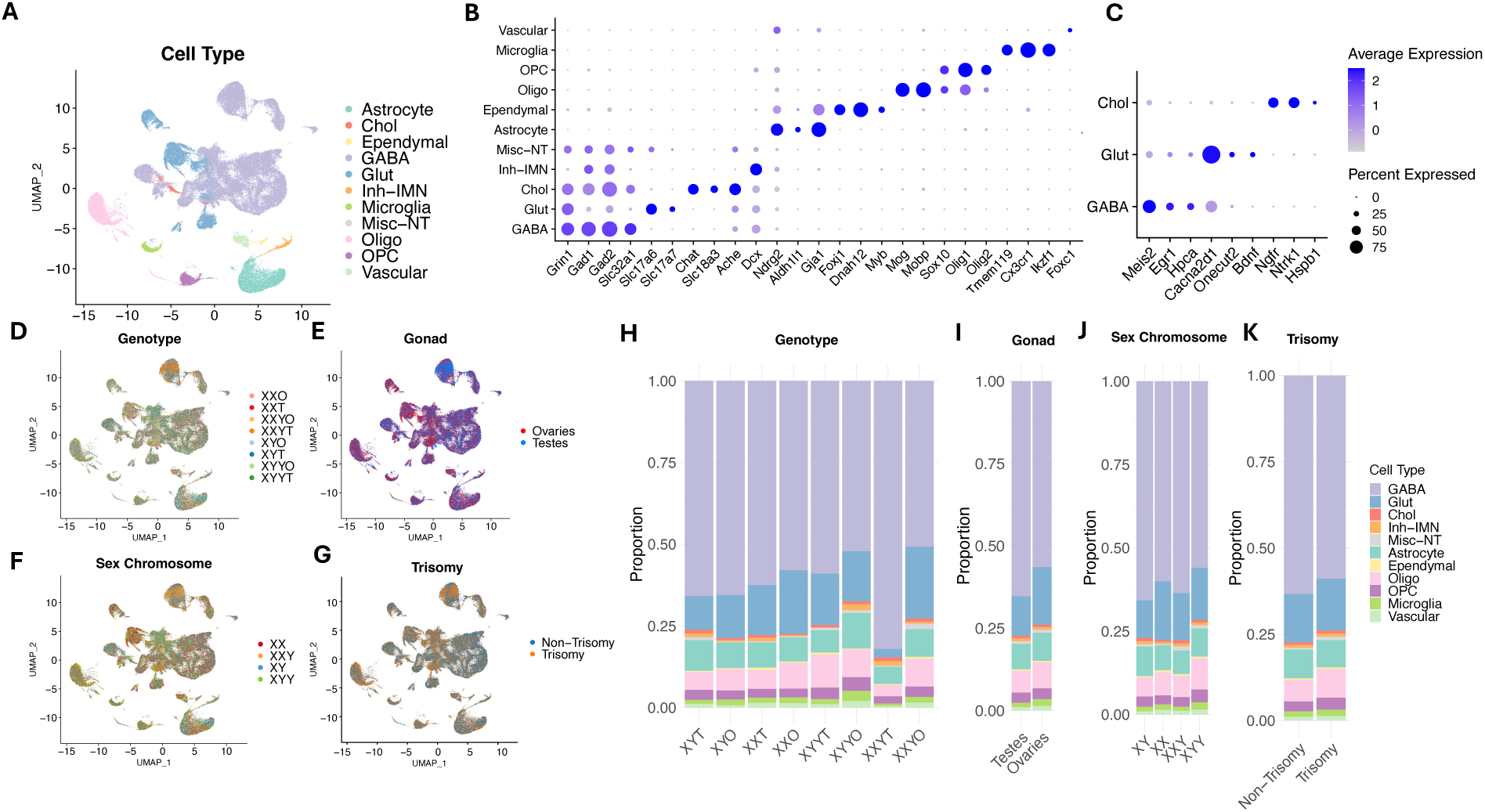
**Diversity of cell types across genotypes and sex factor classifications in the mouse MS** (A) UMAP visualization of MS cell clusters, colored by cell type identity. (B) Dot plot showing scaled expression of canonical marker genes across annotated cell types. Dot size indicates percentage of cells expressing each gene; color represents average scaled expression. (C-F) UMAPs colored by genotype, gonadal sex, sex chromosome complement, and trisomy status. (G) Bar plots showing distribution of counts for cell types across genotypes (H-J) Barplots showing proportion of cell types across gonadal sex, sex chromosome complement, and trisomy status, normalized to total cell counts per group.

We next assessed whether any sex factor drove clustering differences (**Figure 2D-G**). For each cell type, we observe a consistent representation of cells from different genotypes, gonads, sex chromosome complement, and trisomy status, with only a slight XYYT predominance in one GABAergic subcluster.

### Sex-biasing factors do not significantly affect MS cell type proportions

We next assessed whether cell type composition was significantly dissimilar between groups. GABAergic neurons constituted the bulk of cells across all genotypes (**Figure 2H**). However, we found no statistically significant differences in cell type proportions between genotypes or between gonad, sex chromosome, or trisomy status (**Figure 2I-K**) (**Supp Tables 4-7**). Overall, these results suggest sex-biasing factors do not directly modify relative composition of individual cell types.

### Establishing baseline or “typical” sex differences in gene expression across cell types between XXO and XYT

We next defined the “typical” sex differences in gene expression between typically developing XXO and XYT mice using a gene-wise linear model, incorporating genotype as the main explanatory variable and sequencing date as a covariate. To reduce gene expression sparsity, we constructed metacells, which recapitulated the strong clustering and marker gene expression patterns of the major cell type clusters (**Supp Fig 4**) and identified DEGs for each cell type (**Figure 3**). As expected, we observe consistent upregulation of X-linked genes in XXO vs. XYT mice - most significantly *Xist* and *Tsix* - and the expected absence of expression in XXO for several Y-linked genes highly expressed in XYT mice such as *Uty* and *Eif2s3y* across all cell types (**Figure 3A**). Based on the ranking in significant DEG numbers, astrocytes, GABAergic and glutamatergic neurons, microglia, and oligodendrocytes exhibited the most robust typical sex differences (**Figure 3B**). Additionally, most cell types featured a greater number of female-biased DEGs, suggesting that the MS may operate under a female-biased transcriptional program (**Figure 3C**). Finally, the vast majority of the identified DEGs were autosomal (**Figure 3D**), confirming that sex effects are not restricted to genes located on the sex chromosomes.

**Figure 3.**
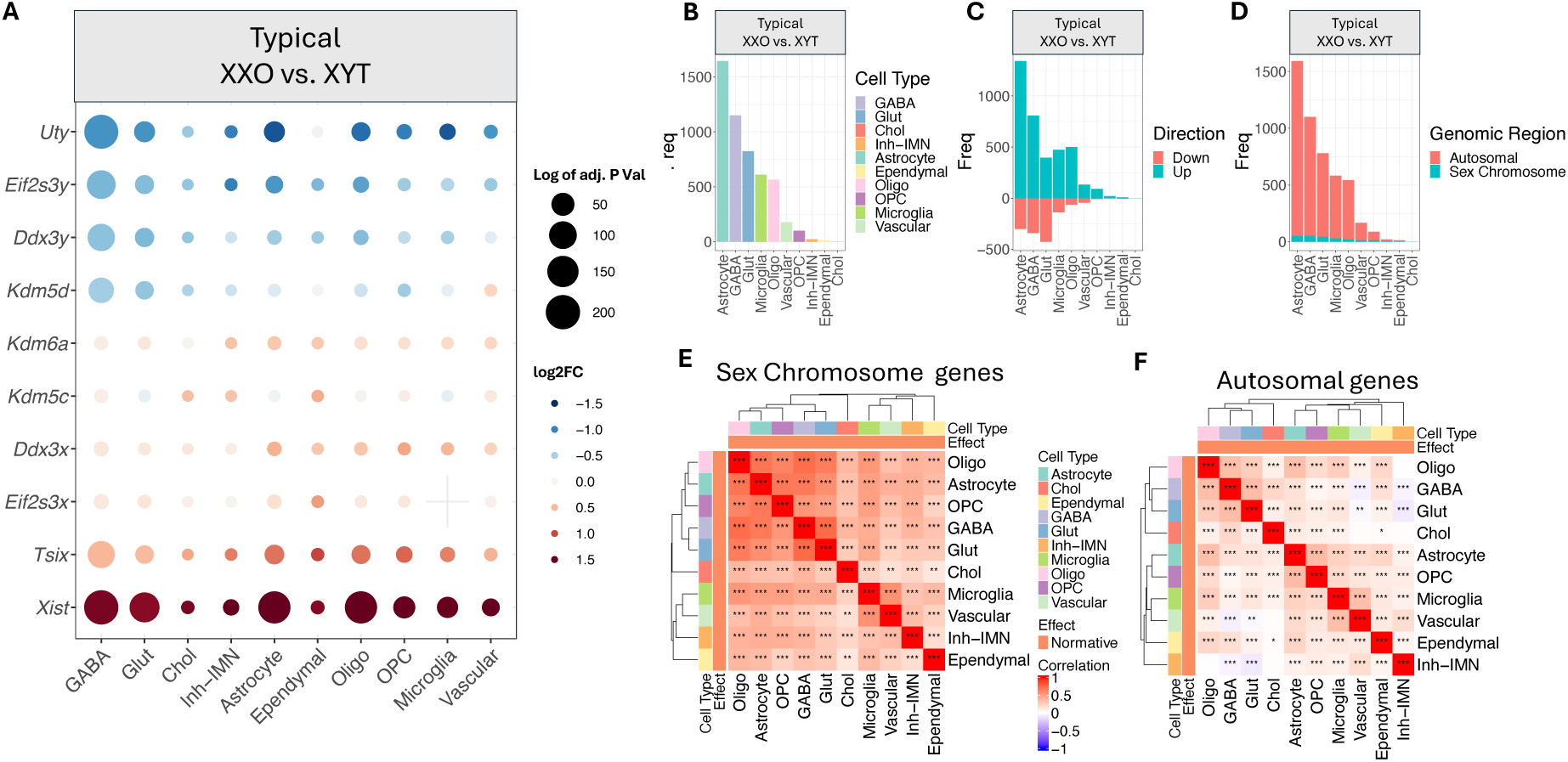
**Cell type-specific typical sex difference DEGs between XXO and XYT in mouse MS** (A) Number of differentially expressed genes (DEGs) for the typical sex difference (XXO vs. XYT) per cell type. (B) Number of differentially expressed genes (DEGs) for the typical sex difference per cell type stratified by direction of change, where “Up” = XXO > XYT, and “Down” = XXO < XYT. (C) Number of differentially expressed genes (DEGs) for the typical difference per cell type colored by the genomic region of the gene. (D-E) Heatmaps displaying pairwise correlations of log2FC values across cell types, calculated separately for (D) sex chromosome genes and (E) autosomal genes using Pearson’s correlation. Dendrograms reflect hierarchical clustering of cell types based on transcriptional correlation. Asterisks denote significance of correlation (* p<0.05, ** p<0.01, *** p<0.001) (F) Dotplot showing DEG result for known gametologues. Dots are colored by the log2FC and sized based on the −log10 transformed adjusted p-value. Positive log2FC (in red) indicates XXO > XYT expression, while negative log2FC (in blue) indicates XXO < XYT.

To further assess commonalities and differences in typical sex differences between cell types, we correlated the log2FC values of the typical sex differences for all genes between cell types, analyzing sex-linked and autosomal genes separately. Given the centrality of sex chromosome dosage differences between XYT and XXO groups, sex chromosome genes display much stronger global correlation patterns across cell types, with groupings driven primarily by cell differentiation lineage (i.e., glial cells and the neuronal subtypes originating from the neuroectoderm and the remaining immune and vascular cells from the mesoderm) (**Figure 3E**). In contrast, autosomal genes display relatively weaker correlations between cell types (**Figure 3F**) and a distinct clustering pattern, with most glial cells separating from neurons. Interestingly, oligodendrocytes consistently clustered closely to neurons, suggesting that these cell types share coregulated sex-biased genes, possibly as part of processes driven by neuron-oligodendrocyte interactions such as myelination.

### Transcriptional response to individual sex factors varies across cell types

Having established the cell type-specific typical sex difference, we next sought to isolate the effects of individual sex-biasing factors using gene-wise linear models (**Figure 4A-C**). As positive controls, we verified the upregulation of *Xist* and *Tsix* and downregulation of *Uty* and *Eif2s3y* for the SCE (XX vs XY) across cell types. We also verified the expected expression patterns for these control genes for the XCD (2 vs 1 X) and the inverse pattern for the YCD1 (1 vs 0 Y) and YCD2 (2 vs 1 Y) effects (**Supp Figure 5**), supporting the accuracy and sensitivity of the FCG and SCT models.

**Figure 4.**
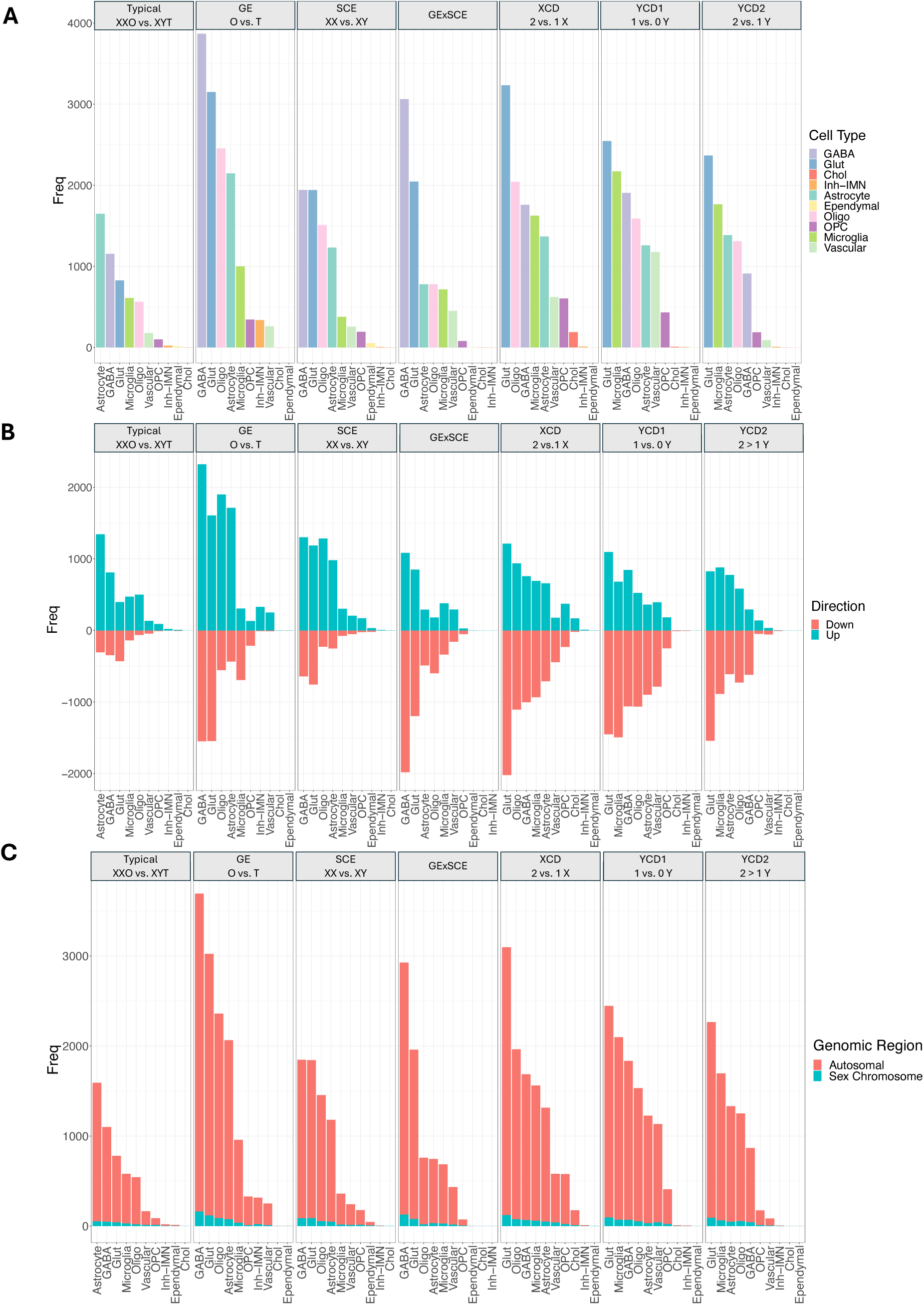
**Cell type-specific transcriptional responses to sex-biasing factors in mouse MS** (A) Bar plots indicating number of significant DEGs across all major MS cell types under seven sex effects and comparisons: normative (“Up” = XXO > XYT, “Down” = XXO < XYT), gonadal sex (“Up” = ovaries > testes, “Down” = ovaries < testes), sex chromosome complement (“Up” = XX > XY, “Down” = XX < XY), interaction of gonad and sex chromosomes, and three trisomy conditions: XCD (“Up” = 2X > 1X, “Down = 2X < 1X), YCD1 (“Up” = 1Y > 0Y, “Down” = 1Y < 0Y), and YCD2 (“Up” = 2Y > 1Y, “Down” = 2Y < 1Y). (B) DEGs stratified by direction of change within each comparison and cell type. (C) DEGs stratified by genomic location within each comparison and cell type.

Analysis of FCG effects (GEs, SCEs, and their interactions, GExSCEs), reveals robust transcriptional responses across a variety of cell types that modulate both autosomal and sex chromosome genes (**Figure 4A,C**). GABAergic and glutamatergic neurons consistently exhibit the largest number of DEGs, in stark contrast to cholinergic neurons, which are resistant to any FCG effect, followed by oligodendrocytes and astrocytes. While our results confirm that sex hormones cause widespread transcriptional changes, we also detect a distinct role of sex chromosomes in regulating gene expression across most cell types (**Figure 4A**). Although GABAergic and glutamatergic neurons exhibit the largest transcriptional response to the SCE, both oligodendrocytes and astrocytes display a comparable number of SCE DEGs. Moreover, many cell types showed an appreciable number of GExSCE DEGs, indicating a distinct transcriptional signature that is dependent on both gonadal hormone exposure and sex chromosome complement. Interestingly, while GABAergic and glutamatergic neurons show a relatively balanced number of female and male-biased GE and SCE DEGs, astrocytes and oligodendrocyte DEGs tend to be female-biased (O > T, XX > XY), reflecting observed patterns in the typical sex difference (**Figure 4B**).

When analyzing sex chromosome dosage effects (XCD, YCD1, YCD2), we observe that glutamatergic neurons consistently exhibit the highest number of DEGs. However, in contrast to FCG effects, glial cells displayed elevated DEG numbers, with oligodendrocytes, microglia, and astrocytes ranking ahead of GABAergic neurons for certain dosage effects. Among glial cells, microglia and astrocytes displayed a consistent transcriptional response, while other cell types were more selective. For example, vascular cells exhibited elevated YCD1 DEGs but few XCD and YCD2 DEGs, suggesting a particular sensitivity to the presence of the first Y chromosome.

In sum, assessment of FCG and SCT effects highlights unique trends across profiled cell types. Importantly, these rankings hold when DEG counts are normalized by the number of expressed genes per cell type (**Supp Figure 6**), indicating that our DEG results are representative of an intrinsic sexually differentiated cellular phenotype rather than reflections of differences in basal gene expression.

### Contributions of individual sex-biasing factors to typical sex differences

Having defined the effects of each sex-biasing factor, we then sought to further dissect the typical sex difference into individual contributing components by quantifying the proportion of typical DEGs overlapping with those of a single sex factor in contrast to those that overlap with multiple (“Shared”) or unexplained by any sex factor (“Typical_Unique”) (**Figure 5**).

**Figure 5.**
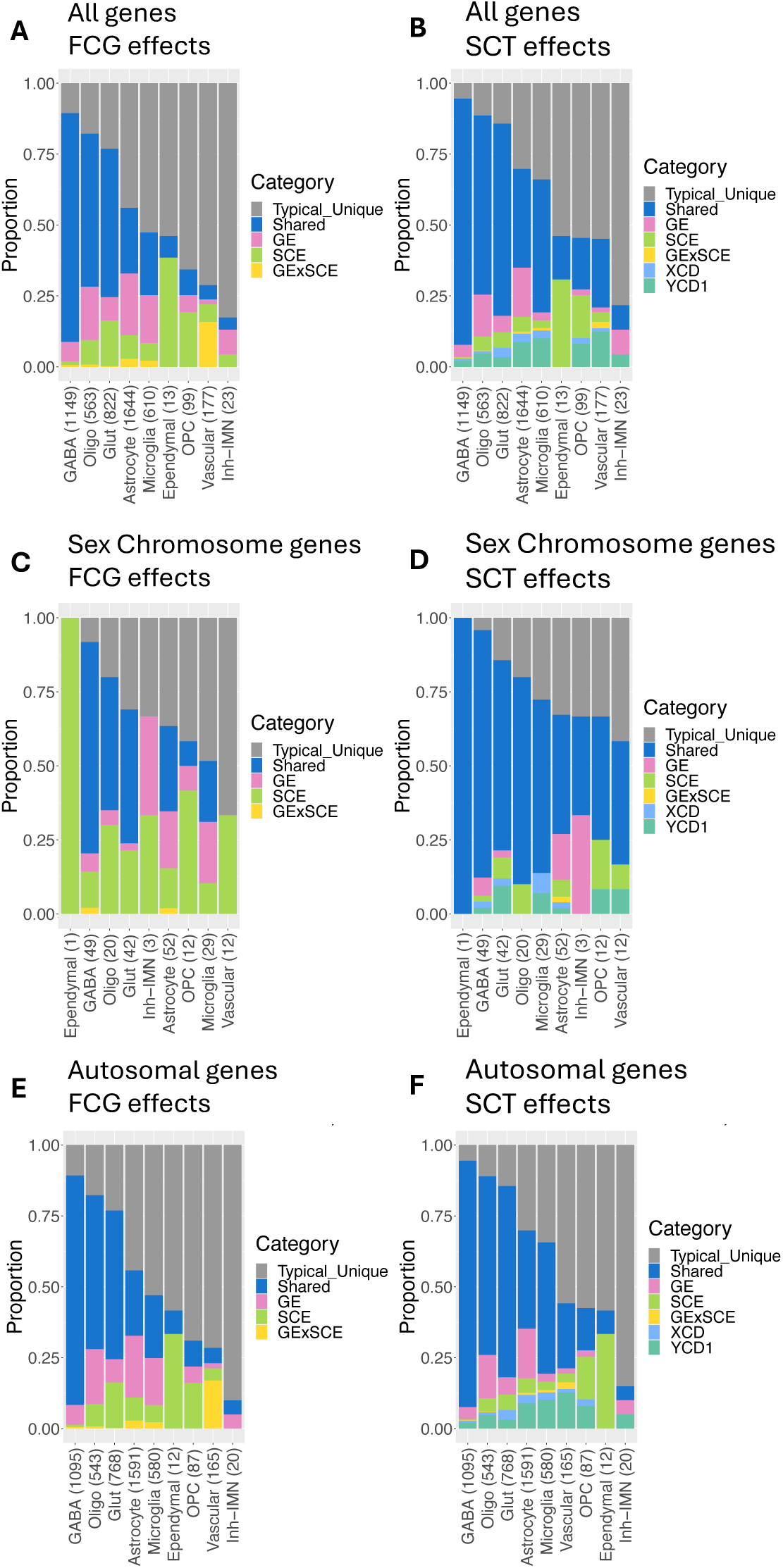
**Correlation of the typical sex difference with distinct sex-biasing factors** (A-D) Bar plots indicating the typical DEG set overlap with DEGs of different sex factors. Results are separated based on genomic location of the gene and the sex factors considered. (A) DEG overlap considering all DEG genes and FCG effects (GE, sex SCE GExSCE). (B) DEG overlap considering all DEG genes and all relevant SCT effects (GE, sex SCE GExSCE, XCD, YCD1). (C) DEG overlap considering sex chromosome genes and FCG effects. (D) DEG overlap considering sex chromosome genes and all relevant SCT effects. (E) DEG overlap considering autosomal genes and FCG effects. (F) DEG overlap considering autosomal genes and all relevant SCT effects.

Results indicate that the degree of the typical sex difference that can be associated with any FCG effect varies by cell type (**Figure 5A**). While >50% of typical DEGs overlap with FCG effects for oligodendrocytes, astrocytes, and GABAergic and glutamatergic neurons, in OPCs, vascular cells, or immature inhibitory neurons the majority of the typical DEGs do not overlap with significant DEGs of any FCG effect. For these cells, their sex differences may arise from cumulative effects of multiple sex-biasing mechanisms each with subtle effects (hence not detected as significant DEGs).

Of the typical sex difference attributable to sex factors, we observe that for GABAergic neurons, oligodendrocytes, and glutamatergic neurons, most DEGs fall into the “Shared” category, indicating that multiple sex factors regulate similar sets of genes in these cell types. However, for others, the largest set is attributable to a single sex factor, with a substantial proportion of sex-biased genes uniquely attributable to SCEs, further stressing their unique contribution in shaping the typical sex difference in most cell types. Correlation and regression of the log2FC values for each sex-biasing factor with the typical effect (see methods) confirms the relevance of sex chromosome-associated effects (**Supp Figure 8**).

By incorporating all SCT effects in this overlap analysis, the SCE can be further decomposed into the unique contributions of the XCD and YCD1 effects. We observed a larger contribution of YCD1 relative to XCD DEGs, indicating that baseline sex differences in gene expression between XXO and XYT cells are driven by the absence of the Y chromosome rather than the added second X chromosome (**Figure 5B**). These observations are recapitulated when separately analyzing sex chromosome DEGs (**Figure 5C,D**) and autosomal genes (**Figure 5E,F**).

### Diverse biological pathways are enriched among sex factor DEGs

To gain insight into the biological functions of the sex-biased DEG sets, we next mapped identified DEGs to pathways sourced from the GO:BP database. To simplify the interpretation of 70 DEG sets (10 cell types x 7 sex effects), we constructed a log2FC correlation matrix across all DEG sets to sort each set into distinct clusters. Consistently DEGs for each cluster were then used as input for pathway overrepresentation analysis (see methods).

Clustering of log2FC correlation patterns allowed for the identification of 8 distinct clusters C1-8 (**Figure 6A**). The majority of clusters were composed of DEG sets associated with specific sex factor(s), and the remainder were defined by cell type identity: C1: neurons with mixed effects; C2: GE and SCE; C3: XCD, YCD1, YCD2; C4: mixed cell types with YCD2 effects; C5: typical; C6: GE, GExSCE; C7: GExSCE; C8: neurons with GExSCE. While correlation patterns within these clusters give insight into similarity between gene sets, patterns between clusters indicate concordance between transcriptional responses of various sex factors. For example, we note that C5 (typical) is positively correlated with C2 (GE, SCE), further emphasizing the relevance of GEs and SCEs in the overall sex difference. Separately, we observe that C5 (typical) is strongly anti-correlated with a subset of C2 that is solely comprised of YCD1 gene sets (outlined in green in **Figure 6A**), corresponding with the expected directional relationship between the YCD1 effect (1 vs 0 Y chromosome) with the typical sex difference (XXO vs XYT).

**Figure 6.**
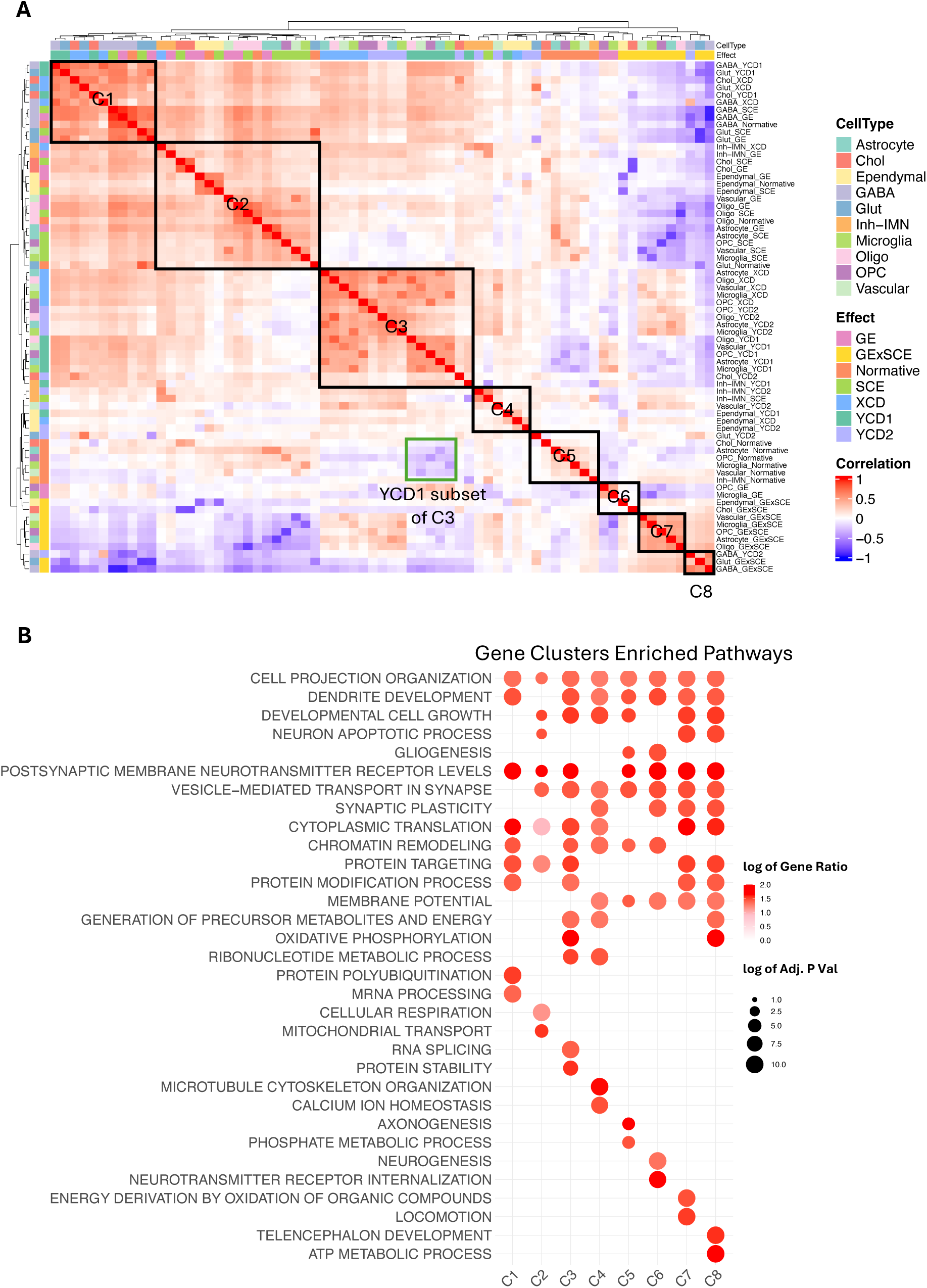
**Transcriptional responses point to clusters of co-expression that map to consistent biological pathways** (A) Heatmaps displaying pairwise correlations of normative log2FC values across cell types and sex effects. Dendrograms reflect hierarchical clustering of cell type and sex effect gene sets based on transcriptional correlation. (B) Over-representation analysis results using GO:BP gene sets and genes from previously identified clusters as input. Color of the dots corresponds to extent of gene set overlap (GeneRatio), while size indicates significance of the −log10 transformed adjusted p-values.

At the pathway level, DEGs across clusters are enriched for a range of processes, including neuronal development, synaptic organization, neurotransmitter secretion and signaling, gene/epigenetic regulation, protein synthesis and modification, and metabolism (**Figure 6B**). Such broad enrichment patterns confirm that a variety of core biological pathways are sex-biased in the MS.

Cluster-specific overrepresented pathways uncover specific effects of individual sex-biasing factors and/or cell types. For instance, C1 (neuronal DEGs with mixed sex factor effects) displays unique enrichment of pathways associated with protein production and degradation, suggesting that the influence of sex factors extends beyond the transcriptomic space to modulate basal neuronal protein expression. While such conclusions have been previously reported, this is the first for the murine MS^53^. In addition, C2, consisting of primarily GE and SCE DEG sets, features pathways centered around cellular metabolism and ATP production, lending further support to the previously reported effects of gonadal hormones and SCEs on mitochondrial metabolism^54,55^. Other unique pathways associated with specific sex factors include RNA splicing and protein stability for C3 (XCD, YCD1, and YCD2), phosphate metabolism, and axonogenesis for C5 (typical).

### Cell type-specific Gene Regulatory Network (GRN) modeling reveals key regulators of sex-biased DEGs in MS

While previous analyses illuminated specific sex-biased genes and biological functions, they did not reveal the broader transcriptional architecture that may provide important clarity into the overall structure of gene programs that dictate cellular responses, facilitating discovery of central and peripheral genes that drive convergence or divergence of distinct sex-biasing effects. To this end, we constructed cell type-specific GRNs (see methods). Once constructed, cell type and sex factor-specific DEG sets were overlaid onto the network, and network hubs whose network neighborhoods were enriched for the DEGs, termed “key drivers” (KDs) were identified based on network topology (see methods). We focused on the glutamatergic neurons for deeper analysis owing to both their strong sex-biased transcriptomic profile and their core functional roles in mediating oscillatory patterns in the MS^56^.

Across the top subnetworks with the strongest statistical significance, both DEGs and KDs are associated with unique combinations of sex-biasing factors, suggesting that combinations of distinct sex factors converge onto and regulate specific transcriptional networks and biological pathways. One subnetwork featured genes associated with synapse maintenance and plasticity, with KDs composed of autosomal genes associated with GE and YCD2 (**Figure 7**). A top key driver was *Homer1*, which encodes a scaffolding protein that regulates glutamatergic synapses and spine morphogenesis. Mutations in this gene are associated with the etiology of a range of neuropsychiatric disorders, including autism spectrum disorder, schizophrenia, and major depression^57,58^. Another key driver was *Ddn*, which encodes the protein Dendrin, which has also been linked to synaptic plasticity^59–61^. Notably, alterations in Dendrin levels are associated with sleep deprivation, suggesting a potential mechanism through which MS glutamatergic neurons, which have been implicated in controlling wakefulness via the septo-hippocampal circuit, may modulate these processes^62^.

**Figure 7.**
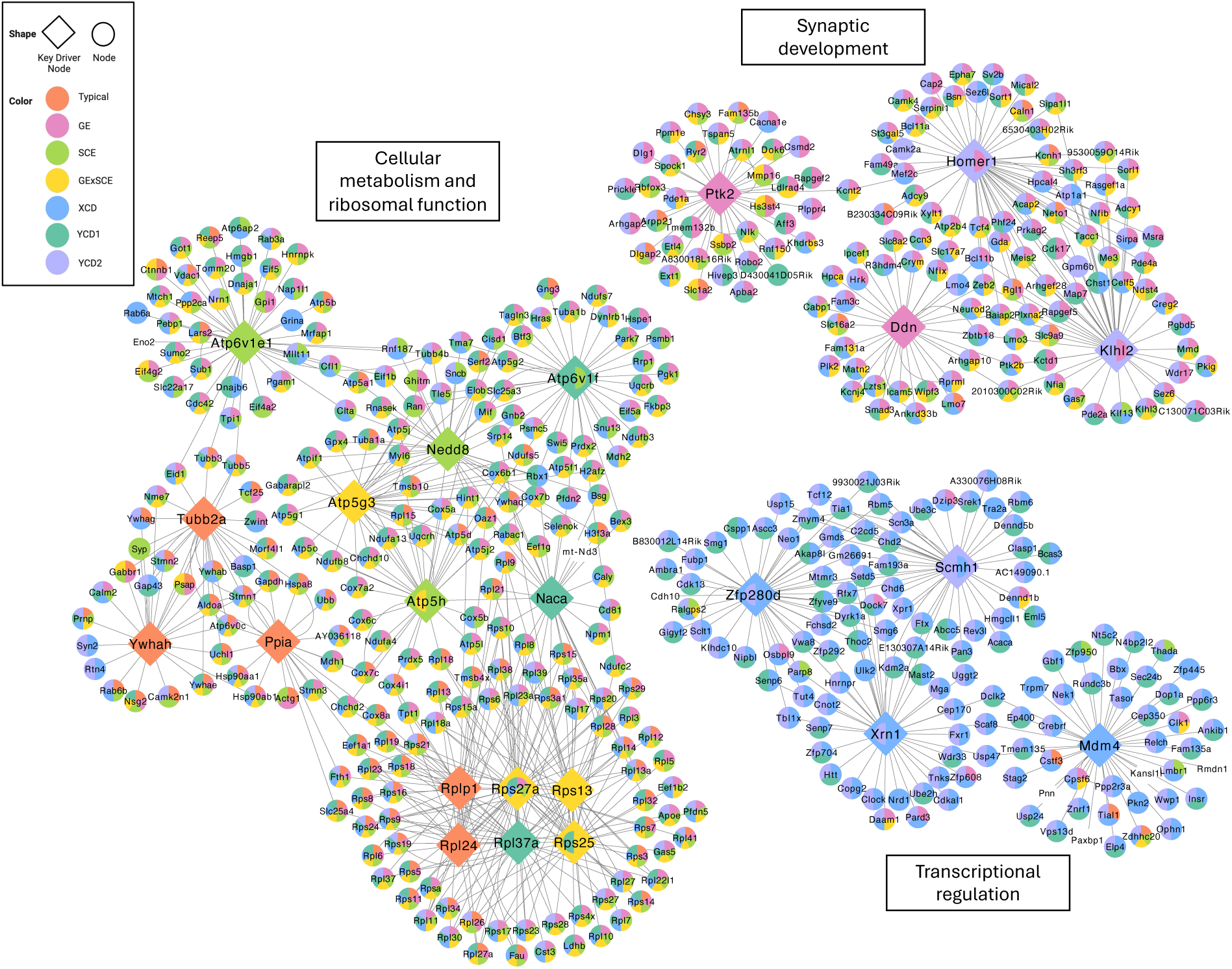
**GRN modeling reveals transcriptional basis of sex differences in Glutamatergic neurons** Visualization of glutamatergic neuron networks around top key drivers identified for sex factor-associated subnetworks. Top key drivers (nodes in diamond shapes) were selected from the top five independent key regulatory genes for each sex factor. Subnetwork member genes denoted as smaller circular nodes and colored by DEG status for each sex factor.

Another identified subnetwork displayed widespread enrichment of KDs associated with transcriptional regulation. For example, *Xrn1* regulates transcriptional dynamics by mediating mRNA degradation, while *Zfp280d* has been predicted to regulate DNA-templated transcription by enabling transcription factor binding activity^63,64^.

The last subnetwork featured KDs associated with the typical sex difference, SCE, GExSCE, and YCD1. Multiple KDs encode subunits of ATP synthase, indicating regulation of energy production and metabolic activity, while the presence of multiple *Rpl* genes points to active protein synthesis and ribosomal activity. This subnetwork strongly indicates active cellular growth and proliferation that is concurrently mediated by multiple sex-biasing factors, particularly SCE-related terms.

### Disease Association of sex factor DEGs

Finally, we used marker set enrichment analysis (MSEA; details in methods) to integrate our findings with human disease genetics, inferring (1) etiologic cell type whose transcriptional profile aligns with human disease genetics and (2) specific sex differentiation mechanisms that may underlie the sex-biased nature of multiple human diseases.

Analysis of 98 human GWAS showed a broad enrichment across traits across cell types and sex factors (**Figure 8**). A range of neurological disorders with known sex biases that have separately been associated with disruptions to theta oscillations were captured through this analysis, including Schizophrenia, Depressive symptoms, Attention Deficit Hyperactive Disorder (ADHD), Aggression, and Depression, via significant enrichments in DEG sets linked to the typical sex difference^65–67^. Although these enrichments implicate sex-biased DEGs from a range of cell types, they most consistently are associated with GABAergic neurons and astrocytes, two cell types that have previously been identified as active contributors to a range of neuropsychiatric disorders^68,69^.

**Figure 8.**
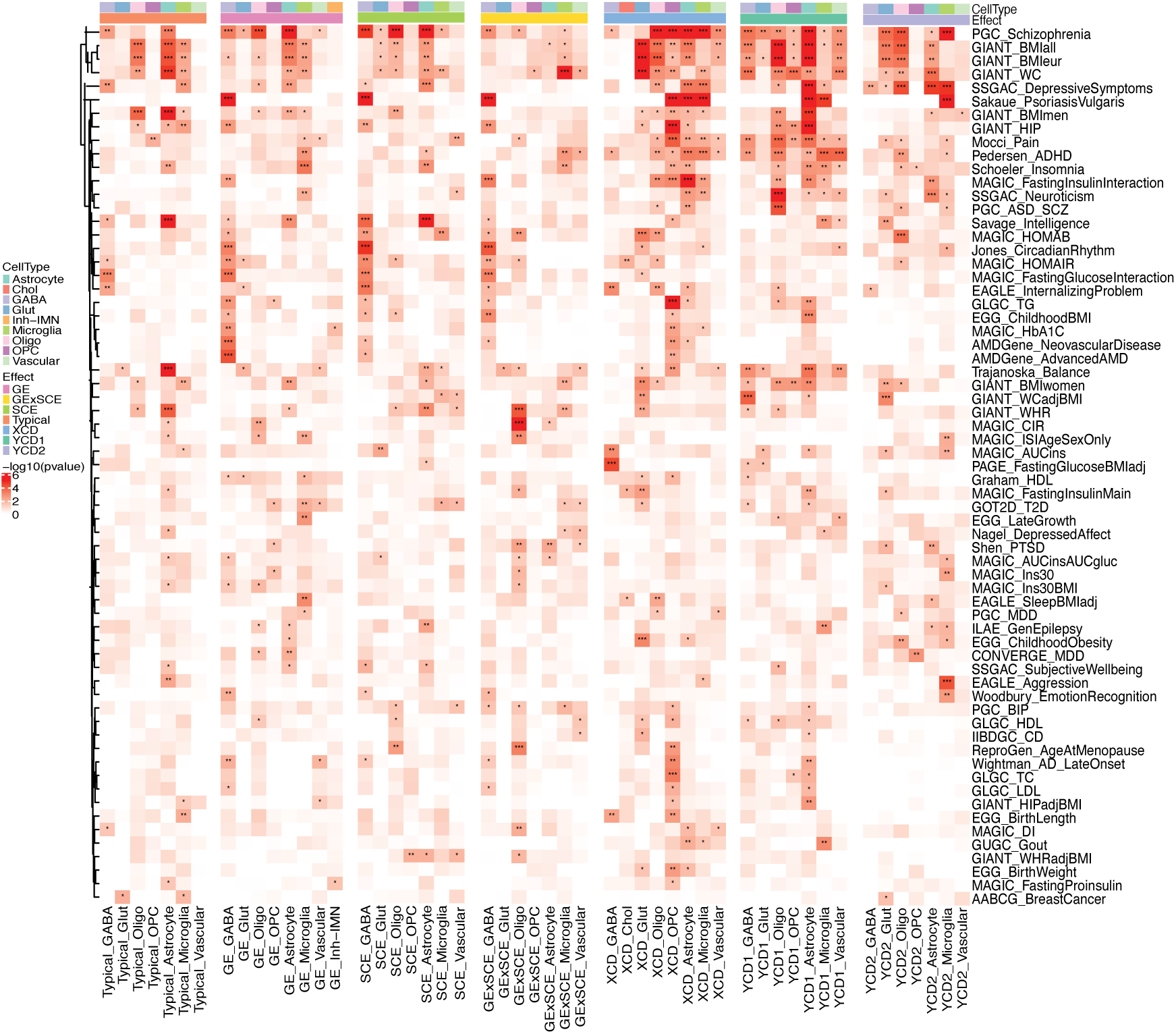
Enrichment of DEGs with genes associated with human disease. Heatmap showing results of MSEA. Cell type-specific DEGs across sex effects used as input alongside mapped genes from human GWAS summary statistics targeting a range of metabolic, neuropsychiatric, and cognitive traits. Cells clustered by sex effect and cell type identity. Color corresponds the degree of significance of enrichment based on −log10 transformed adjusted p values, with asterisks denoting specific significance thresholds (* p<0.05, ** p<0.01, *** p<0.001).

Closer inspection of FCG and SCT effects further sheds light on specific mechanisms that potentially underlie the sex differences in disease genetics. For example, we observe a broad and robust enrichment of sex-biased DEGs across most detected cell types and sex effects for human Schizophrenia GWAS signals. This pattern suggests that the MS is an etiologic region contributing to sex differences in schizophrenia, likely through intercellular signaling dynamics governed by multiple interacting sex-biasing factors across a range of cell types. These findings are consistent with previous studies that have established the role of MS in modulating psychosis-related behaviors and consciousness^70,70,71^.

Similar analysis also gives insight into patterns implicating specific cell types and sex factors in neuropsychiatric disorders that have not been widely associated with the MS. For example, for ADHD, we observe the strongest enrichment in gene sets driven by the XCD and YCD1 effects in glial cells, suggesting that changes in X and Y chromosome dosage rather than sex hormone levels, may drive the disparate patterns in disease risk between females and males in humans. These results recapitulate the known role of disturbed glial cell function in ADHD while also pinpointing sex-chromosome-regulated genes as the specific sites of dysregulation, aligning with prior studies linking X chromosome absence with increased ADHD symptom severity, including attention problems, hyperactivity, and lack of inhibitory control^72^.

Although XCD and YCD1 have been used thus far to dissect the SCE, they can also be used to infer the cell type-basis of the increased disease risk for individuals with trisomy syndromes such as XYY syndrome and Kleinfelter’s syndrome. For example, individuals with XYY syndrome are known to be susceptible to a range of health issues, including learning disabilities, developmental delay, ADHD, and Autism Spectrum Disorder (ASD)^73–76^. Inspection of YCD2 gene sets show that microglia and oligodendrocytes show significant enrichment with ASD as well as ADHD, suggesting that the effects from the second Y chromosome in these cell types may play a role in this increased disease risk. The pathological roles of oligodendrocytes and microglia in ASD have been previously identified in the mouse amygdala, and our findings indicate that similar dynamics may play in the MS in XYY individuals^77^.

Finally, because the MS is not known to directly regulate systemic metabolism, these diseases/traits were included as potential negative controls. However, we unexpectedly identified many such traits and disorders to be strongly enriched among the sex-biased DEG sets across MS cell types, suggesting a potential novel function of MS.

## Discussion

Substantial and reproducible sex differences are observed across diverse domains of human health and disease, including well-documented male biases in early-emerging cognitive impairments and female biases in adolescent-emergent mood and neurological disorders^78^. These stereotyped patterns suggest sex differences in brain function and development. While sex differences in brain anatomy have been thoroughly explored, cellular and molecular-level studies that profile sex differences in gene expression and their key modulators are lacking. To address this, our study provides a more comprehensive understanding than previous studies of not only the overall sex difference in gene expression, but also its underlying sex-biasing mechanisms on a cellular basis in the mouse MS, a region critical for memory consolidation, learning, and hippocampal theta rhythmogenesis. Our findings suggest that the bulk of cell types in the MS operate in a sex-differentiated manner and are sensitive to multiple independent sex-biasing factors. In contrast to the classical, gonad-centric model of sexual differentiation, our analysis points to the high relevance of sex chromosome-associated effects in the overall typical sex difference as well as in sex-biased disease genetics. This work offers crucial insight into the functional basis of sex differences in a poorly studied brain region, with far-reaching implications for understanding the mechanisms driving the sex-biased nature of cognitive and neuropsychiatric disorders.

Using the SCT mouse model, we demonstrate that the MS exhibits robust typical sex differences in gene expression (**Figure 3**). In line with numerous previous studies that have identified sex differences in neuronal and glial cell function, morphology, gene expression, and activity^79–82^, we observe a distinct transcriptional signature in response to the typical sex difference that encompasses a broad range of cell types, including GABAergic and glutamatergic neurons, astrocytes, and oligodendrocytes. Such sex-specific behavior has been previously documented in sexually dimorphic brain regions such as the hippocampus as well as in *in vitro* culture systems^83^, but our study is the first to do so in the MS.

The utility of the SCT model lies in its ability to not only profile typical sex differences, but also to parse the effects of X- and Y-linked sex-biasing factors. Assessment of FCG and SCT effects reveals distinct patterns between cell types in the relative contribution of each sex-biasing factor to gene expression and regulation (**Figure 4**). We observed robust GEs affecting GABAergic and glutamatergic neurons, oligodendrocytes, astrocytes, and to a lesser extent, microglia. Given the well-documented role of sex hormones in modulating diverse neuronal and glial cellular physiology, these results are unsurprising^80,84–86^.

However, our findings reveal extensive SCEs and GExSCEs across a comparable range of cell types. Similarly, we observe a substantial number of sex chromosome dosage-related DEGs associated with XCD, YCD1, and YCD2 effects. Comparing DEG patterns across sex factors and the concordance of their direction of change provides compelling evidence that distinct sex factors interact within cell types. For example, in microglia, the majority of typical and SCE DEGs are female-biased (XXO > XYT; XX > XY), whereas most GE DEGs are male-biased (T > O). These distributions suggest that the influences of sex chromosomes may interact with and counterbalance the influence of gonads, effectively nullifying the opposing GE in the overall typical sex difference^87,88^. Similarly, microglia displayed higher numbers of DEGs for sex chromosome dosage effects relative to SCEs, suggesting potential compensatory mechanisms, where XCD and YCD influences on gene expression may interact and negate each other’s effects such that transcriptional patterns due to each sex factor are not reflected in the cumulative SCE effect. Such patterns showcase the utility of the SCT model in its ability to reveal counterbalancing sex effects that are not reflected in typical female vs male comparisons.

DEG overlap analysis further emphasized the heightened relevance of SCEs and YCD1 in the typical sex difference, compared to GEs, suggesting that the presence of the Y chromosome and its genes can be associated with a non-trivial proportion of the typical sex difference in the mouse MS (**Figure 5**). Such findings are striking as traditionally, the Y chromosome has been thought to primarily mediate differentiation and function of the testes. However, recent studies have increasingly focused on its role in health and disease, linking its presence to higher risks of male-biased diseases such as autism^89^. In sum, our results highlight the importance of sex chromosome-associated effects in driving the sex-biased gene regulation of MS cell types, corroborating previous studies that have suggested certain sex differences in brain phenotype and gene expression are independent of gonadal hormone action in the mouse brain^4,90^.

Interestingly, distinct sex-biasing factors converge and feature similarity in their DEG sets and functional pathways to a degree. Shared pathways across DEGs of sex-biasing factors are involved in synaptic signaling, plasticity, development, and organization, cell growth, protein translation and modification, and cellular metabolism (**Figure 6**). This functional overlap supports previous findings that sex-biased genes often belong to pathways associated with energy production, protein synthesis and transport, among others^91^. We also further identified unique pathways driven by specific sex-biasing factors, such as mitochondrial transport and cellular respiration driven by C2 (GEs, SCEs), or protein and mRNA processing for C1 (neuronal).

Given the extensive transcriptomic responses to sex-biasing factors observed across MS cell types, we further explored their potential regulators by employing a GRN modeling approach. We identified three distinct submodules within glutamatergic neurons that govern diverse processes, namely transcriptional control, synaptic development, and cellular metabolism, along with their key regulators such as *Homer1* and *Xrn1* (**Figure 7**). Because these genes may play a significant role in mediating fundamental biological functions in a sex-biased manner, they serve as intriguing targets for future validation studies. Of note, the constructed GRNs exhibit a high degree of connectivity and complex organization, with genes that are often affected by multiple sex-biasing factors. Such trends suggest that distinct sex-biasing factors can coalesce onto the same regulatory nodes and transcriptional programs in order to govern core cellular phenotypes, highlighting the far-reaching scope of sex-biasing factors in regulating cellular phenotypes.

Taken together, converging lines of evidence across various analyses suggest that the sex-biased anatomical variation of the MS is also reflected in the sex-biased cellular gene expression profiles that may alter cellular function. Crucially, the direction of observed anatomical variation is largely consistent with the direction of gene expression changes across cell types, with a persistent pattern of wild type female > male as well as XX > XY sex biased gene expression. Via pathway enrichment and GRN modeling, we show that these genes consistently regulate biological functions associated with cellular growth, energy production, and synaptic development and pruning, which may underlie observed female-biased anatomical patterns. Such emphasis on SCEs independent of gonadal hormones has been previously identified in the mouse hippocampus, suggesting that the septo-hippocampal circuit as a whole may be primarily mediated by SCEs^92^.

Finally, to further evaluate the potential connections of the sex-biasing genes from our mouse MS study to human disease as a means to parse the mechanisms driving sex-biased disease risks, we connected our findings to a range of available GWAS summary statistics (**Figure 8**). The results provide molecular support for the established role of the MS in psychosis, with strong enrichments for sex-biased diseases such as ADHD and schizophrenia and implicating specifically GABAergic neurons and astrocytes as potential etiologic cell types. Comparison of trends between sex-biasing factors indicate that relative to GEs, dosage effects display more consistent enrichment across cell types. This emphasis on dosage effects recapitulates prior findings that have identified how they modulate features such as brain anatomy, neuronal gene expression, and development^93–96^.

Intriguingly, our MSEA findings extend beyond neuropsychiatric and cognitive disorders, revealing multiple signals associated with metabolic phenotypes, such as BMI and waist circumference, which have previously been shown to be strongly affected by sex-biasing factors^97^. Given that a causal role for the MS in regulating physiological traits has not been established, we propose two potential explanations for this enrichment. First, this pattern may be due to the direct connection of the MS with the hippocampus, an etiologic brain region for obesity whose contribution to food-intake and metabolism regulation has been increasingly recognized^98,99^. While the lateral septum has been known to mediate hippocampal-hypothalamus signaling to modulate glucose concentrations, a potential role of the MS may be suggested here^100^. Conversely, this finding could suggest that even though sex factors induce similar molecular and transcriptional profiles in cell types across brain regions, it is only in the correct molecular context that a cell type is truly causal for human disease. These two possibilities emphasize the poorly understood nature of the MS and reinforces the need for future research to determine its precise contribution to human health.

## Limitations

Although our findings present interesting implications for the MS, we also acknowledge the limitations of our study. First, we note that the SCT model consists of mice from both disomic and trisomic backgrounds, with the SCE representing the difference caused by the contrast of mice with two sex chromosomes (XX vs. XY), that includes effects of X and Y dosage (1X vs. 2X and 0Y vs. 1Y). However, the XCD and YCD1/2 factors measure the effect of an additional X/Y chromosome in the context of shifting from the disomic to trisomic state. Thus, we do not expect *a priori* that dosage effects reported in this study will completely mirror dosage differences in a disomic context. To address this, future studies will consider additional incorporation of samples from the XY* model, which includes mice with XO, XX, XY, and XXY genotypes^101^. While similar to the SCT model, the XY* model can also directly distinguish effects caused by differences in X chromosome dosage (XO vs. XX) from Y chromosome dosage (XO vs. XY) within a disomic state, which may more accurately reflect baseline sex differences (XXO vs. XYT)^102–104^.

Another limitation of our study is that the GEs identified throughout cell types could originate from two separate mechanisms. First, GEs could arise due to an established sex-biased neuronal circuitry generated during key developmental periods, or alternatively, as the product of a transient response to gonadal hormone exposure prior to tissue extraction. These contrasting potential causal mechanisms are rooted in the separation between “organizational” (long-lasting) versus “activational” (reversible) gonadal effects, which can only be distinguished by comparing physiological responses before and after gonadal hormone administration in gonadectomized samples^105^. The current analysis does not distinguish between these mechanisms but may offer an interesting direction for future research.

## Data availability

snRNAseq data is being deposited to Gene Expression Omnibus and accession number is pending.

## Code availability

Code used in this study is available at https://github.com/JShin97/medial_septum_sct or by

contacting the lead author.

## Declarations

### Ethics Approvals

All experiments were approved by the Institutional Animal Care and Use Committees of participating institutions.

### Consent for publication

Not applicable.

## Acknowledgements

This work was supported by the NIH-NICHD grant R01HD100298 (A.P.A, A.M.G, J.L, A.R, J.S, X.Y), NIH-NIMH grant R21MH129020 (X.Y), and the NIH Training Grant in Genomic Analysis and Interpretation T32HG002536 (J.S). The authors thank the University of California, Los Angeles Technology Center for Genomics and Bioinformatics for sequencing of the snRNA-seq data.

## Author Contributions

A.P.A, A.M.G, J.L, A.R, J.S, X.Y conceived and designed the project. H.H and A.M.G managed mouse breeding and dissected tissue samples. G.D, I.S.A, G.Z performed scRNA-seq. J.S, C.M performed bioinformatics analyses and generated figures. J.S, A.P.A, A.R, A.M.G, J.L, M.B, X.Y drafted and revised the manuscript.

