## Supplemental Figures for "Relative contribution of gonads and sex chromosomes to sex differences in cell-type gene expression in the mouse medial septum and sex-biased disease risk"

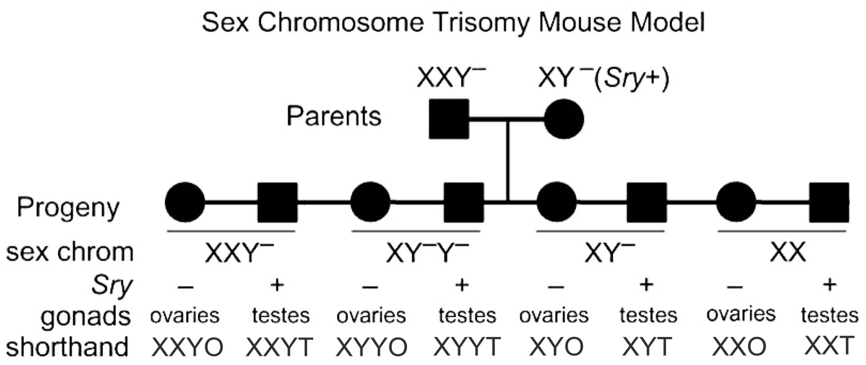


**Supplemental Figure 1. Schematic of the Sex Chromosome Trisomy (SCT) mouse model**

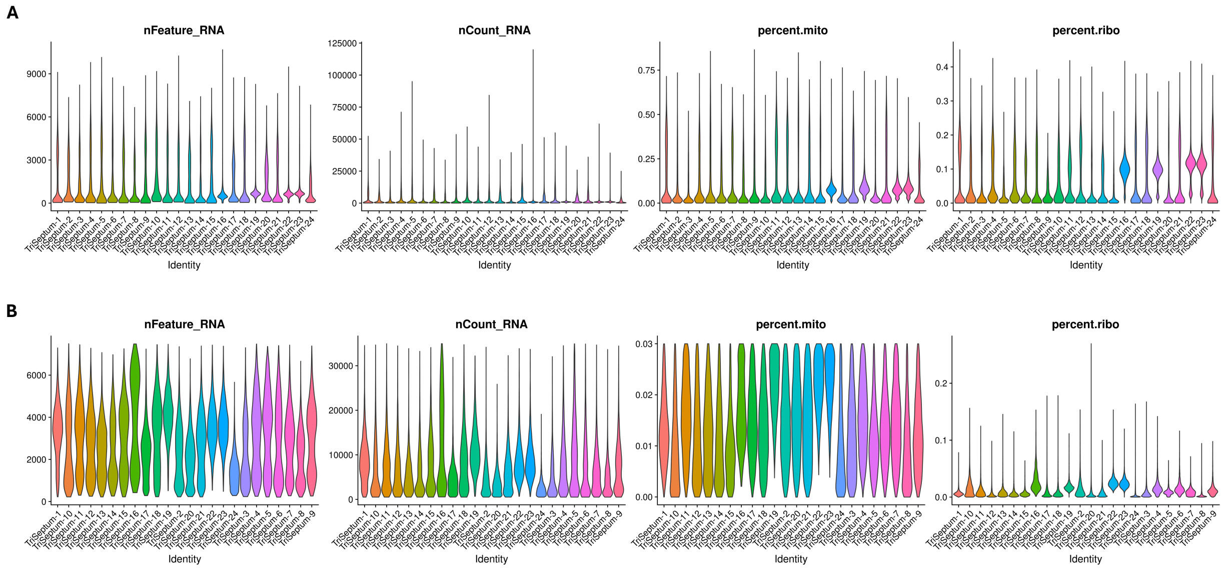


**Supplemental Figure 2. Sample-wise metrics pre- and post-QC**

(A,B) Distribution of number of unique genes detected (nFeature_RNA), total number of UMIs (nCount_RNA), percentage of mitochondrial genes (percent.mito), and percentage of ribosomal genes (percent.ribo) before (A) and after (B) QC.

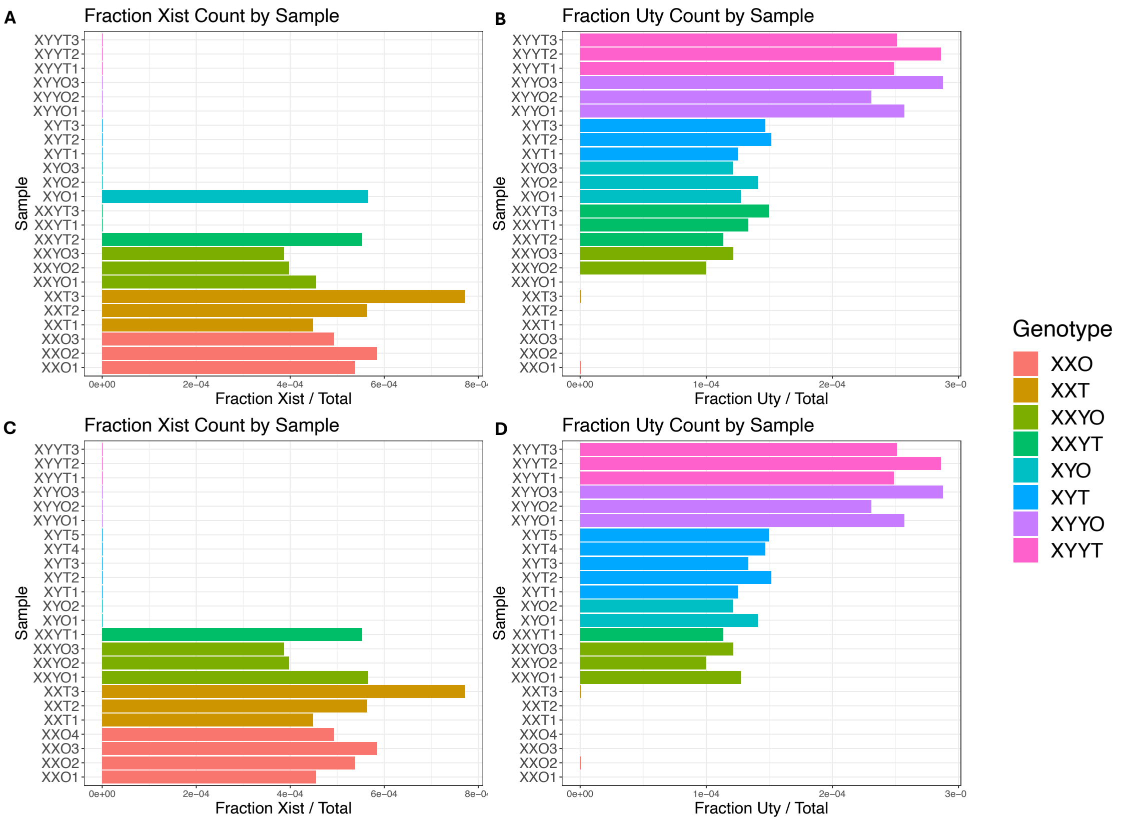


**Supplemental Figure 3. Sample-wise gene counts for *Xist* and *Uty affirms revised genotyping.***

(A-B) Fraction of gene counts for (B) Xist and (C) Uty across sequenced samples, with color of bars corresponding to the original assigned genotype.

(C-D) Fraction of gene counts for (B) Xist and (C) Uty across sequenced samples, with color of bars corresponding to the updated genotype. Reassignment corresponds to expected patterns of Xist and Uty expression by genotype.

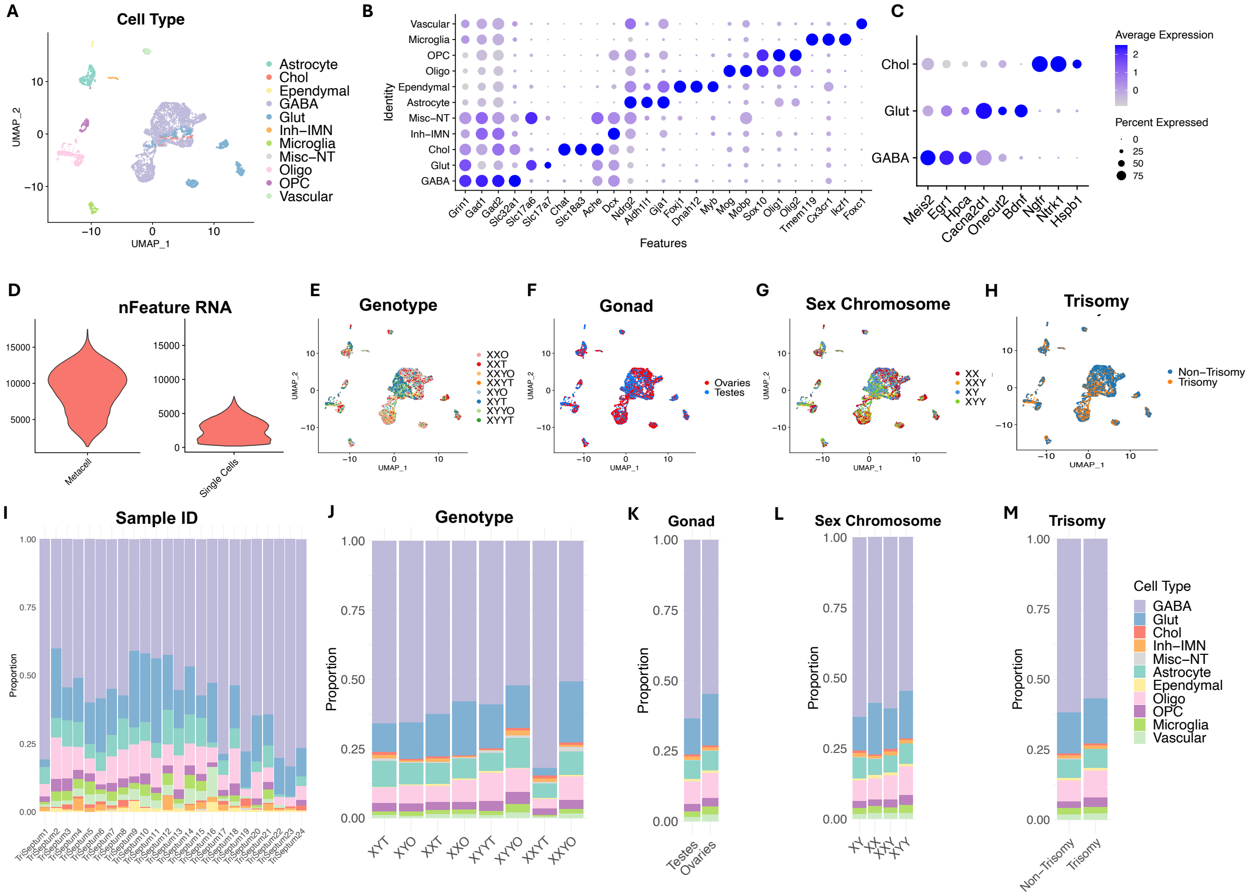


**Supplemental Figure 4: Metacells reduce data sparsity while preserving biological signal across MS cell types**

A) UMAP visualization of MS metacell clusters, colored by cell type identity.

(B) Dot plot showing scaled expression of canonical marker genes across annotated cell types. Dot size indicates percentage of cells expressing each gene; color represents average scaled expression.

(C) Dot plot showing scaled expression of MS-specific neuronal marker genes across annotated cell types. Dot size indicates percentage of cells expressing each gene; color represents average scaled expression.

(D) Violin plot of the number of detected genes for metacells (left) compared to single cells (right).

(E-H) UMAPs of metacells colored by genotype, gonadal sex, sex chromosome complement, and trisomy status

(I-M) Barplot showing metacell counts when grouped by sample ID, genotype, gonadal sex, sex chromosome complement, and trisomy status.

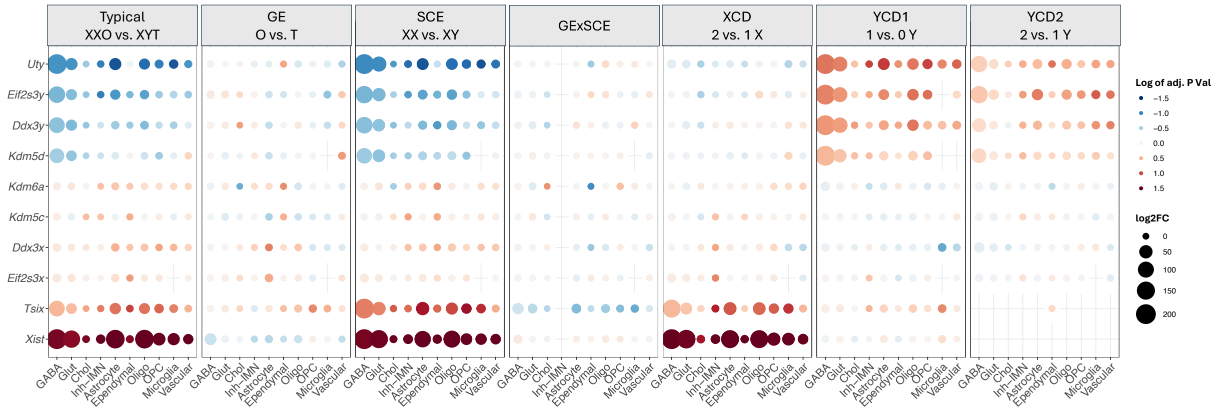


**Supplemental Figure 5: Sex-linked genes are captured in relevant DEG models using SCT mice.** log2FC was calculated as the female to male effect for the typical sex difference and FCG effects and as an increasing dosage effect for XCD and YCD1/2 terms: typical - XXO vs XYT; GE - O vs T; SCE - XX vs XY; XCD - 2 vs 1 X; YCD1 - 1 vs 0 Y; YCD2 - 2 vs 1 Y. Red and blue indicate higher and lower in the first group, respectively.

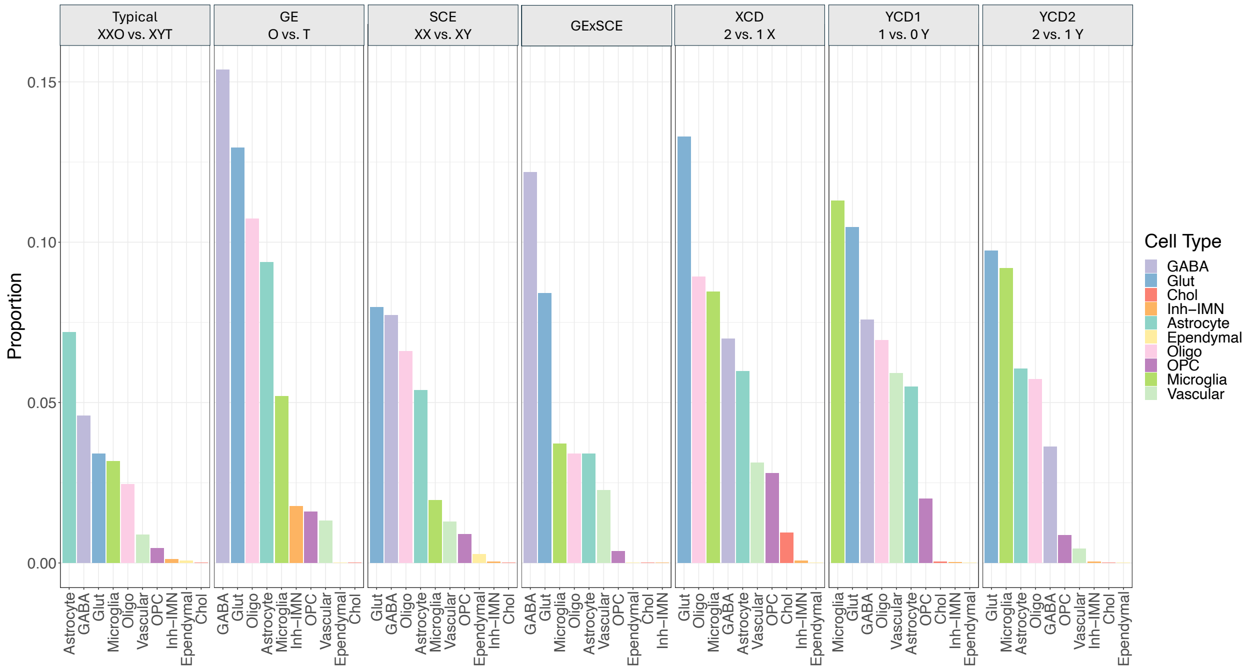


**Supplemental Figure 6: Sex-biasing DEGs normalized by number of expressed genes**

The proportion of the significant DEG number (p.adjust < 0.05 and abs(log2FC) > 0.1) normalized by the number of expressed genes within each cell type is plotted across all cell type and gene effect combinations.

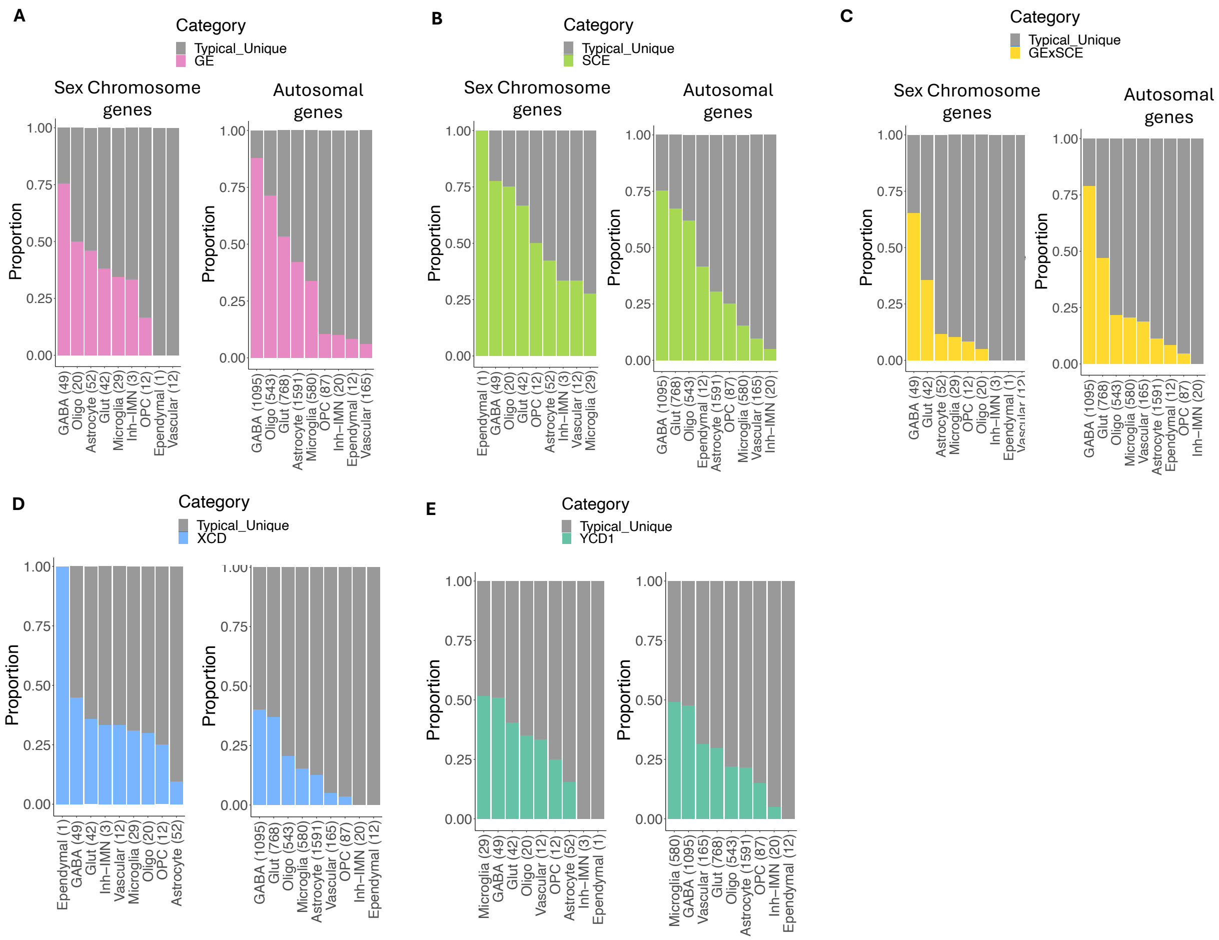


**Supplemental Figure 7: Gene overlap results considering 1 sex effect at a time**

(A-E) The typical sex difference was associated with individual sex-biasing effects based on gene set overlap. Results are shown for when considering

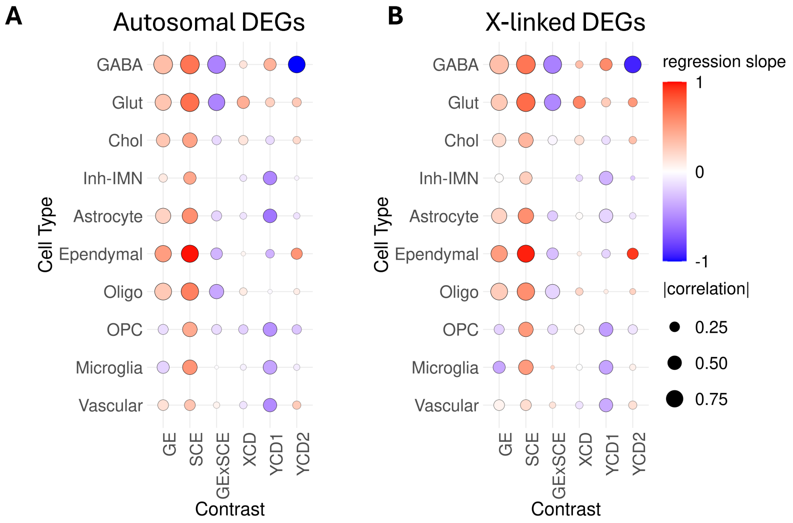


**Supplemental Figure 8: log2FC correlation and regression results across cell types and sex effects.**

(A-B) Heatmaps indicating the similarity of log2FC between the typical sex difference in gene expression and the effects of all sex-biasing effects assessed by the SCT and FCG models (columns) for all cell types (rows). Color of the dot corresponds to the regression slope and size indicates correlation strength. Results shown separately for autosomal (A) and X-chromosome genes (B).

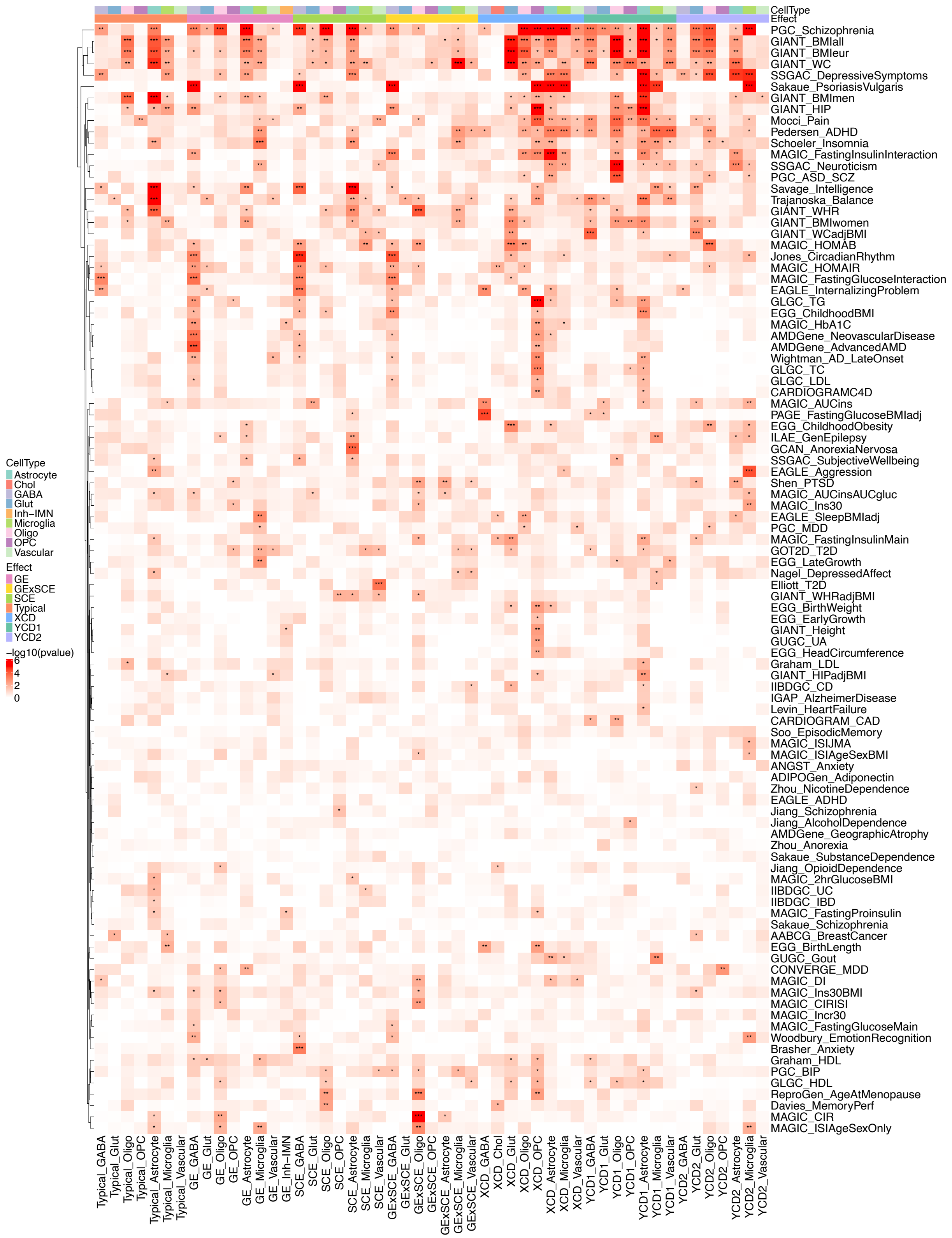


**Supplemental Figure 9: MSEA results for all datasets**

Heatmap showing unfiltered MSEA results. Cell type-specific DEGs across sex effects used as input alongside mapped genes from available human GWAS summary statistics targeting a range of metabolic, neuropsychiatric, and cognitive traits. Cells clustered by sex effect and cell type identity. Color corresponds the degree of significance of enrichment based on -log10 transformed adjusted p values, with asterisks denoting specific significance thresholds (* $p<0.05$, ** $p<0.01$, *** $p<0.001$).
